# Heterogeneity in quiescent Müller glia in the uninjured zebrafish retina drive differential responses following photoreceptor ablation

**DOI:** 10.1101/2023.01.27.525802

**Authors:** Aaron J. Krylov, Shuguang Yu, Axel Newton, Jie He, Patricia R. Jusuf

## Abstract

Loss of neurons in the neural retina is a leading cause of vision loss. While humans do not possess the capacity for retinal regeneration, zebrafish can achieve this through activation of resident Müller glia. Remarkably, despite the presence of Müller glia in humans and other mammalian vertebrates, these cells lack an intrinsic ability to contribute to regeneration. Upon activation, zebrafish Müller glia can adopt a stem cell-like state, undergo proliferation and generate new neurons. However, the underlying molecular mechanisms of this activation subsequent retinal regeneration remains unclear. To address this, we performed single-cell RNA sequencing (scRNA-seq) and report remarkable heterogeneity in gene expression within quiescent Müller glia across distinct dorsal, central and ventral retina pools of such cells. Next, we utilised a genetically driven, chemically inducible nitroreductase approach to study Müller glia activation following selective ablation of three distinct photoreceptor subtypes: long wavelength sensitive cones, short wavelength sensitive cones, and rods. There, our data revealed that a region-specific bias in activation of Müller glia exists in the zebrafish retina, and this is independent of the distribution of the ablated cell type across retinal regions. Notably, gene ontology analysis revealed that injury-responsive dorsal and central Müller glia express genes related to dorsal/ventral pattern formation, growth factor activity, and regulation of developmental process. Through scRNA-seq analysis, we identify a shared genetic program underlying initial Müller glia activation and cell cycle entry, followed by differences that drive the fate of regenerating neurons. We observed an initial expression of AP-1 and injury-responsive transcription factors, followed by genes involved in Notch signalling, ribosome biogenesis and gliogenesis, and finally expression of cell cycle, chromatin remodeling and microtubule-associated genes. Taken together, our findings document the regional specificity of gene expression within quiescent Müller glia and demonstrate unique Müller glia activation and regeneration features following neural ablation. These findings will improve our understanding of the molecular pathways relevant to neural regeneration in the retina.

## 1 Introduction

Vision loss results from dysfunction or death of retinal neurons, and this condition represents a significant global health issue that affects billions of people worldwide (Stevens et al., 2013; Bourne et al., 2017). Irreversible vision loss because of retinal neuron death can be caused by genetic factors, environmental insult as well as age-dependent decline (Gordois et al., 2012). One attractive approach to restore functional vision is to harness the endogenous capacity for neuronal regeneration within the retina.

Unlike humans, zebrafish possess an intrinsic ability to regenerate the retina following injury (Nagashima et al., 2013; Lenkowski and Raymond, 2014) so as to restore visual function (Sherpa et al., 2008). The regenerative capacity of zebrafish is dependent on retinal glial cells, known as Müller glia. During retinal regeneration, Müller glia are capable of cellular de-differentiation through a myriad of genetic (Fausett et al., 2008; Ramachandran et al., 2010; Ramachandran et al., 2012; Gorsuch and Hyde, 2014; Elsaeidi et al., 2018; Thomas et al., 2018; Sharma et al., 2020) and epigenetic changes (Powell et al., 2013; Mitra et al., 2018; Hoang et al., 2020) so as to adopt a stem cell-like state. Subsequent cell proliferation leads to production of clonal progenitors (Vihtelic and Hyde, 2000; Fausett and Goldman, 2006; Bernardos et al., 2007). These progenitors can migrate and differentiate into neurons within sites of tissue damage (Montgomery et al., 2010; Powell et al., 2016; Ng Chi Kei et al., 2017; D’Orazi et al., 2020; Lahne et al., 2021).

Despite mammalian Müller glia showing evidence of cell cycle entry in response to retinal injury (Dyer and Cepko, 2000; Yoshida et al., 2004; Kase et al., 2006; Conedera et al., 2021), these cells do not replace lost neurons and instead form scarring at the site of injury, termed reactive gliosis (Wilhelmsson et al., 2004; Bringmann et al., 2006). Across vertebrates, the gene expression profiles for Müller glia are remarkably conserved (Bringmann et al., 2009a; Thomas et al., 2016), with neuroprotective (Harada et al., 2002; Hauck et al., 2006; Bringmann and Wiedemann, 2012; Furuya et al., 2012; Reichenbach and Bringmann, 2013) and homeostatic roles (Newman et al., 1984; Yamada et al., 1999; Bringmann et al., 2009b; MacDonald et al., 2015) conserved in vertebrate species. Despite such conservation in gene expression and function, there is currently no evidence that Müller glia can generate new neurons within the human. Nevertheless, it is recognised that genes such as *ascl1a* (Fausett et al., 2008), as well as members of the Wnt (Ramachandran et al., 2011), Sonic hedgehog (Thomas et al., 2018) and Hippo signalling pathways (Lourenço et al., 2021) are important mediators of the Müller glia proliferative response in zebrafish. These genes have led to improvements in the neurogenic ability of mammalian Müller glia (Pollak et al., 2013; Ueki et al., 2015; Todd et al., 2016; Yao et al., 2016; Jorstad et al., 2017; Yao et al., 2018; Hamon et al., 2019; Rueda et al., 2019; Jorstad et al., 2020; Todd et al., 2021). Thus, by improving our understanding of Müller glia-driven regeneration in zebrafish, we can better understand neurogenesis in mammalian Müller glia, including in humans.

The gene networks that underlie Müller glia-driven neurogenic responses in vertebrates remains poorly understood. Indeed, while it is known that a subset of Müller glia proliferate following injury (Bailey et al., 2010; Thomas et al., 2016), it is not clear if the regenerative capacity for Müller glia across the entire retina is shared, or if there are region-specific or injury-specific features of such a response. Here, we addressed such issues through a series of single-cell RNA sequencing (scRNA-seq) studies using Müller glia isolated from Tg(*gfap:GFP*) zebrafish through fluorescent activated cell sorting (FACS), whereby GFP expression specifically in mature Müller glia of the retina is under the control of a gene promoter for glial fibrillary acidic protein (GFAP). Through this approach, we describe distinct subpopulations of quiescent and/or resting Müller glia that are regionally distributed across the retina, along the dorsal ventral axis. Further, to determine the response to injury, we generated zebrafish lines to model the loss of specific photoreceptor subtypes and found that distinct subpopulations of Müller glia mount unique regenerative responses. Additionally, we characterised the gene expression modules of Müller glia from quiescence to activation, where we show evidence for three cellular states of Müller glia following photoreceptor injury. Taken together, this study enhances our understanding of the genetic drivers and potential barriers of retinal regeneration in zebrafish that could support our understanding of the potential for human Müller glia to be used to treat neuron loss.

## 2 Methods and Materials

### 2.0 Zebrafish husbandry

All zebrafish strains were bred and housed in the Danio *rerio* research facilities within the University of Melbourne and the Walter and Eliza Hall Institute of Medical (ethics approval ID 22235 and 10400), in accordance with local guidelines. All procedures were approved by the Faculty of Science Animal Ethics Committee at the University of Melbourne (approval IDs 1814542 and 10232). Zebrafish embryos were raised at 28.5°C in E3 medium (5 mM NaCl, 0.17 mM KCl, 0.33 mM CaCl2, 0.33 mM MgSO4) at a maximum density of n = 50 per 50 ml petri dish. E3 medium was supplemented with 0.003% 1-phenyl-2-thiourea (PTU) from 24 hours post-fertilisation (hpf) to prevent pigmentation and to maintain transparency of zebrafish embryos, prior to sorting for fluorescence signal. Prior to sorting for experiments, transgenic Tg(*gfap:GFP*) larvae were anaesthetised in 0.04 % tricaine methanesulfonate (MS-222; Sigma-Aldrich, cat. number E10521-50G) for > 24 hours hpf, while larvae for Tg(*lws2:nfsb-mCherry*), Tg(*xops:nfsb-mCherry*) and Tg(*sws2:nfsb-mCherry*) strains were treated with the same bath for > 72 hpf.

### 2.1 Generation of transgenic lines

DNA constructs were injected into one-cell stage wildtype AB zebrafish embryos, using a FemtoJet microinjector (Eppendorf) and borosilicate glass capillary needle (1.0 mm O.D. / 0.78 mm I.D. / 100 mm long capillary). Tg(*gfap:GFP*) zebrafish were generated by Bernardos and Raymond, 2006. lws2:nfsb-mCherry, xops:nfsb-mCherry and sws2:nfsb-mCherry DNA constructs were microinjected with tol2 RNA. A pTol-uas:nfsB-mCherry plasmid (a gift from Dr. Toshio Ohshima (Waseda University, Tokyo, Japan) was used to generate these three constructs (Shimizu et al., 2015; Yoshimatsu et al., 2016). The promoters were chosen to specifically ablate the subtype expressing the distinct opsin protein, which is a hallmark of the different photoreceptor types. The sws2:nfsb-mcherry construct was a gift from Prof. Rachel Wong (University of Washington, Seattle, Washington, USA). Plasmid linearisation of Tol2 DNA was achieved using a Quiaquick Kit, while RNA synthesis was carried out using the Ambion message machine kit Sp6.

### 2.2 Single-cell sample preparation

Zebrafish lines Tg(*her4.1:dRFP/gfap:GFP*) was used for control (non-ablated) tissue, while Tg(*her4.1:dRFP/gfap:GFP/lws2:nfsb-mCherry*) and Tg(*her4.1:dRFP/gfap:GFP/sws2:nfsb-mCherry*) were used to obtain long wavelength sensitive (Lws2) cone-ablated and short wavelength sensitive (Sws2) cone-ablated retinal tissue respectively. Zebrafish larvae at 6 days post-fertilisation (dpf) were exposed to a 10 mM solution of metronidazole for 48 hours, to specifically eliminate the long or short wavelength sensitive cone photoreceptor subtype. At 8 dpf, fish were rinsed in fresh system water and housed in standard conditions. At 9 dpf (3 days post-injury; dpi), fish were humanely killed. Single-cell suspensions of 9 dpf zebrafish were prepared using a specific protocol (Lopez-Ramirez et al., 2016). Retinae were dissected and digested in 350 μl papain solution at 37°C for 15 minutes. The papain solution was prepared as follows: 100 μl papain (Worthington, LS003126), 100 ul of 1 % DNAse (Sigma, DN25), and 200 μl of 12mg / ml L-cysteine (Sigma, C6852) were added to a 5 ml DMEM/F12 (Invitrogen, 11330032). During digestion, retinal tissue was mixed by repeated pipetting 4 to 10 times. Following digestion, 1400 μl of washing buffer was added, containing 65 μl of 45 % glucose (Invitrogen, 04196545 SB), 50 ul of 1M HEPES (Sigma, H4034), and 500 μl FBS (Gibco, 10270106) in 9.385 ml of 1x DPBS (Invitrogen, 14190-144). All solutions were filtered through a 0.22 μm filter (Millipore) and stored at 4°C prior to use.

### 2.3 10X Chromium single-cell RNA sequencing

To perform single-cell RNA sequencing (scRNA-seq), cells isolated through fluorescent activated cell sorting (FACS) were loaded onto a Chromium Single Cell Chip (10x Genomics, USA) according to the manufacturer’s protocol. The scRNA-seq libraries were generated using the GemCode Single-Cell Instrument and Single Cell 3’ Library and Gel Bead kit v2 and v3 Chip kit (10x Genomics, 120237). Library quantification and quality assessments were performed by Qubit fluorometric assay (Invitrogen) and dsDNA High Sensitivity Assay Kit (AATI, DNF-474-0500). Analysis of DNA fragments was performed using the High Sensitivity Large Fragment −50kb Analysis Kit (AATI, DNF-464). The indexed library was tested for quality and sequenced using an Illumina NovaSeq 6000 sequencer with the S2 flow cell using paired-end 150 base pair reads. Sequencing depth was 60K reads per cell.

### 2.4 Quality filtering and pre-processing

Filtered matrix raw data files were analysed in R using Seurat. Prior to downstream filtering, the following describes cell numbers for datasets: 8325 (no ablation control), 10520 (Lws2-ablation) and 7851 (Sws2-ablation). Post-filtering and exclusion of non-Müller glia-derived cell types achieved the following cell numbers: 7884 (no ablation control), 9789 (Lws2-ablation), and 7542 (Sws2-ablation). Low quality cells or cells containing doublets were excluded from all datasets; reads between 200 and 3500 genes per cell (no ablation control), reads between 200 and 4500 genes per cell (Lws2-ablation), and reads between 200 and 4000 genes per cell (Sws2-ablation) were obtained. Cells with a percentage of mitochondrial gene expression of greater than 20 % (no ablation control), 35 % (Lws2-ablation), and 30 % (Sws2-ablation) were excluded from further analysis.

Samples were analysed using Seurat::NormalizeData, while variable features for downstream analysis were identified using Seurat::FindVariableFeatures, and scaled using Seurat::ScaleData. An optimal number of principal components (PCs), generated through Seurat::RunPCA, for dimensional reduction were selected using the function Seurat::ElbowPlot. PCs containing the greatest variance were selected. For each sample, 20 PCs were specified, and we selected 13 (no ablation control), 15 (Lws2-ablation) and 15 (Sws2-ablation) PCs. Clustering was performed using the shared nearest neighbour (SNN), graph-based approach through Seurat::FindNeighbours. Uniform manifold approximation and projection (UMAP) plots containing 11 (no ablation control), 12 (Lws2-ablation) and 12 (Sws2-ablation) clusters were prepared using Seurat::FindClusters.

### 2.5 Integration analysis

Samples were integrated by a standard integration protocol for R package Seurat using Seurat::FindIntegrationAnchors and Seurat::IntegrateData. Downstream analysis including data scaling, dimensional reduction and clustering was conducted as per above. A total of 13 PCs (no ablation control vs Lws2-ablation) and 15 PCs (Lws2-ablation vs Sws2-ablation) were selected for dimensional reduction.

### 2.6 Cluster gene expression analysis

Markers driving the characterisation of each cluster were resolved using Seurat::FindMarkers, with only positive features expressed in at least 25 % of cells in each group of cells. Lists of differentially expressed genes generated through this approach were used for gene set enrichment analysis (GSEA), conducted using gprofiler::gost, with statistical significance evaluated using a Benjamini-Hochberg FDR test set at a threshold of 0.05. Relevant terms were selected and plotted in accordance to their negative adjusted *p* value, along with gene ratio (number of genes attributed to the relevant GO term / number of genes in the cluster gene list). Genes mentioned from this cluster characterisation analysis were within the top 50 markers.

### 2.7 Trajectory analysis

Pseudotime analysis was conducted using the R package monocle3. Prior to analysis of trajectory and searching for differentially expressed genes, ribosomal protein genes or mitochondrial genes were identified from lists and excluded. The Seurat object was converted to a cell data set (cds), and a subset of cells from quiescent to activated Müller glia cell states was selected using Monocle3::choose_graph_segments and Monocle3::choose_cells. Next, Monocle3::order_cells was used to select the root node, which was in the quiescent cell cluster furthest from the activated cell population in the UMAP plot. In Monocle3::graph_test, we used the principal graph option with Moran’s test statistic to identify significantly differentially expressed genes along the specified trajectory. The q value was set to < 0.05 and Moran’s I value set to > 0.25. The top 40 differentially expressed genes are presented in a heatmap, hierarchically clustered based on gene modules generated from Monocle::graph_test.

### 2.8 RNAscope in situ hybridisation

RNAscope probes for *efnb2a, fgf24*, and *rdh10a* were generated by ACD company (Shanghai). The experiments were carried out as follows. Larval zebrafish were fixed in 4% PFA at 4°C overnight followed by cryoprotection in 30% sucrose and then were cryosectioned at a thickness of 12 μm. The slices were post-fixed in 4% PFA at room temperature for 15 mins and washed with 1 × PBS at room temperature for 3 mins. To block the activity of endogenous peroxidase, all slides were treated with 0.1 % H2O2 at room temperature for 30 mins. After being washed twice with 1 × PBS at room temperature for 3 mins, slides were treated with 10 μg / ml proteinase K (Sigma) diluted in TE (10 mM Tris-HCl, pH 8.0, and 1 mM EDTA, pH 8.0) at 37°C for 8 mins, then treated with 4 % PFA at room temperature for 10 mins. Subsequently, all slides were washed with 1 × PBS at RT for 3 mins, followed by the incubation in 0.2 M HCl at RT for 10 mins. After washing with 1 × PBS for 5 mins, all slices were then incubated with 0.1 M triethanol amine-HCl (662.5 μl triethanolamine and 1.35 ml 1 M HCl; adding water to the final volume of 50 ml, pH 8.0) at room temperature for 1 min and in 0.1 M triethanol amine-HCl containing 0.25% acetic anhydrate at room temperature for 10 mins with gentle shaking. Slides were then washed by 1 × PBS at room temperature for 5 mins then dehydrated in a series of 60%, 80%, 95% ethanol baths, and finally twice in 100% ethanol at room temperature for 90 seconds, respectively. Slides were incubated in the hybridization buffer (50% formamide (Sigma), 10 mM Tris-HCl, pH 8.0, 200 μg / ml yeast tRNA (Invitrogen), 1 × Denhart buffer, SDS, EDTA and 10 % dextran sulfate (Ambion) containing 1 μg / ml probes at 60°C overnight. On the second day, slides were washed sequentially in 5 × SSC at 65°C for 30 mins, 2 × SSC with 50 % formamide at 65°C for 30 mins, TNE buffer (100 ml TNE consisting of 1 ml 1 M Tris-HCl, pH 7.5, 10 ml 5 M NaCl, and 0.2 ml 0.5 M EDTA) at 37°C for 10 min and then in TNE buffer with 20 μg / ml RNaseA at 37°C for 30 mins. Slides were then incubated with 2 × SSC at 60°C for 20 mins, 0.2 × SSC at 60°C for 20 mins, and 0.1 × SSC at RT for 20 mins. Next, slides were blocked by TN buffer at room temperature for 5 mins (200 ml TN buffer consisting of 20 ml 1 M Tris-HCl, pH 7.5, 6 ml 5 M NaCl, and 174 ml water) followed by TNB buffer (TN buffer and 0.5% blocking reagent; Roche) at room temperature for 5 mins. Finally, slides were incubated in TNB buffer with anti–DIG-POD (1:500; Roche) at 4°C overnight. On the third day, the signal was detected by the TSATM Plus Cyanine 3 / Fluorescein System (PerkinElmer, NEL753001KT).

### 2.9 Metronidazole treatment

For Tg(*lws2:nfsb-mCherry*), Tg(*sws2:nfsb-mCherry*), or Tg(*xops:nfsb-mCherry*) zebrafish larvae processed for immunohistochemical analysis, metronidazole treatment was conducted at 4 dpf for 48 hours, leading to ablation of either Lws2 cones, Sws2 cones, or rod photoreceptors respectively. Zebrafish larvae were exposed to a solution of 10 mM metronidazole (Sigma-Aldrich, cat. number M3761-100G) in standard fish water. To induce a lesser degree of injury in Tg(*lws2:nfsb-mCherry*) zebrafish, larvae were swum in a solution of 5 mM metronidazole in standard fish water for 2 hours. Zebrafish larvae were placed into solution at maximum densities of n = 50 larvae per 50 ml petri dish and kept at 28.5°C for the duration of treatment. Control larvae were placed in standard fish water. Zebrafish were rinsed in standard fish water after treatment. For adult zebrafish, these were immersed in a solution of 10 mM metronidazole in standard fish water for 3 consecutive days, while control adult zebrafish were placed in standard fish water, both with these respective baths changed twice a day.

### 2.10 Histological processing

Zebrafish larvae were humanely killed in 4 g / L tricaine, fixed in 4% paraformaldehyde (PFA; Sigma-Aldrich, cat. number 158127-500G) in PBS overnight at 4°C and subsequently rinsed in PBS. Larvae were then cryoprotected in a 30% sucrose solution in 1x PBS overnight at 4°C before being embedded in OCT (Tissue-Tek) in 10 mm / 10 mm / 5 mm Tissue-Tek Cryomolds (ProSciTech). Moulds containing these tissues were frozen and stored at −20°C. Samples were sectioned at 12 μm thickness using a Microtome Blade (Arthur Bailey Surgico) on a Leica (CM1860) cryostat, with sections transferred onto room-temperature 25 mm / 75 mm Menzel Superfrost Plus glass slides (Grale Scientific), allowed to dry for 1 hour, and used immediately for immunohistochemistry or stored at - 20°C.

### 2.11 Immunohistochemistry

Antibody staining was carried out at room temperature using standard protocols. Antigen retrieval to detect epitopes for antibodies was performed by incubating slides in 150 mM Tris-HCl (pH 9) at 70°C for 20 minutes, then rinsed for 30 minutes before overnight incubation in a primary antibody prepared with 5% foetal bovine serum (FBS) blocking solution. Primary antibodies used for this study were as follows: mouse anti-proliferating cell nuclear antigen (PCNA; Santa Cruz Biotechnology, sc-25290), 1:500; rabbit anti-mCherry (Invitrogen, PA5-34974), 1:500, rabbit anti-Bmpr1b (GeneTex, GTX128200). The next day, slides were rinsed in 1 x PBS and then incubated for 2 hours in secondary antibody, diluted in the same blocking solution. Secondary antibodies used were as follows: Alexa Fluor 647 nm donkey anti-mouse, 1:500; Alexa Fluor 546 nm goat anti-rabbit, 1:500. Following antibody staining, slides were rinsed thrice with 1 x PBS and then stained with 4’,6-diamidino-2-phenylindole (DAPI; 1:10000, Sigma-Aldrich, D9542-10MG) in 1 x PBS solution. Finally, sections were mounted in Mowiol (Sigma-Aldrich, cat. number 81381-250G) using 24 mm / 60 mm glass coverslips (ProSciTech). Slides were stored at 4°C.

### 2.12 Image acquisition and analysis

Images of immunostained immobilized sections were captured using a Nikon A1 confocal microscope with a 40 x pan-fluor oil-immersion objective lens, with 1.3 numerical aperture. Z-stacks were acquired with a step size of 1 μm. For RNAscope samples, images were taken using an inverted confocal microscope system (FV1200, Olympus) using a 30 x (silicon oil, 1.05 NA) or 60 x (silicon oil, 1.3 NA) objective lens. For each fish, an image of a single retina was obtained. Quantification was conducted using FIJI/ImageJ. To define dorsal, central and ventral regions of the retina, three segments of 60° were partitioned for quantification studies therein.

### 2.13 Statistical analysis

Data are expressed as mean ±standard error. The number of fish in each condition are specified in the respective graphs. Statistical analyses were conducted using Prism 8 (GraphPad) using a two-way ANOVA followed by a Bonferroni post hoc test for multiple comparisons. To determine significance between mCherry-positive glia in the activated population compared to the total glia analysed, a chi-square test of independence was conducted. Statistical significance are as follows: * = p ≤ .05, ** = p ≤ .01, *** = p ≤ .001, or **** = p ≤ .0001.

### 2.14 Figure creation

Figures were compiled using Adobe Illustrator. Microscope images were arranged in Adobe Photoshop, where brightness was adjusted evenly across channels and images, prior to placement into Illustrator.

## 3 Results

### 3.0 Müller glia follow a shared quiescence to activation pathway between photoreceptor ablation paradigms

Following neural ablation or cell death in the zebrafish retina, Müller glia that proliferate must first undergo reprogramming from a quiescent to an activated state. First, we characterised the gene expression networks which underlie the switch from quiescence to activation following two injury paradigms; specific ablation of long wavelength sensitive cone photoreceptors (Lws2) in Tg(*lws2:nfsb-mCherry*), or short wavelength sensitive cone photoreceptors (Sws2) in Tg(*sws2:nfsb-mCherry*) zebrafish. Expression of nitroreductase (nfsb) in these cell populations specifically leads to cytotoxicity upon metronidazole exposure (Curado et al., 2007; Curado et al., 2008). We combined our injury lines with Tg(*gfap:GFP*) zebrafish and performed single cell RNA sequencing (scRNA-seq) of fluorescent activated cell sorting (FACS) Gfap-positive cells, 3 days post injury (dpi) in larval zebrafish. By integrating scRNA-seq datasets of Müller glia from Lws2-ablated and Sws2-ablated retinas, we sought to determine commonalities and differences in the Müller glia response (Figure 1). Regardless of injury, quiescent and proliferative clusters were conserved across both datasets. However the main differences were observed in clusters containing differentiating Müller glia-derived progenitor cells (Figure 1A, B). Understanding the molecular pathways that guide the activation of stem cells is of great interest for studying regeneration. To investigate this, we isolated quiescent and activated Müller glia clusters and constructed a pseudotime trajectory (Figure 1C, D) to determine the activation of gene modules which accompany the changes between these two cellular states (Figure 1C – G).

**Figure 1:**
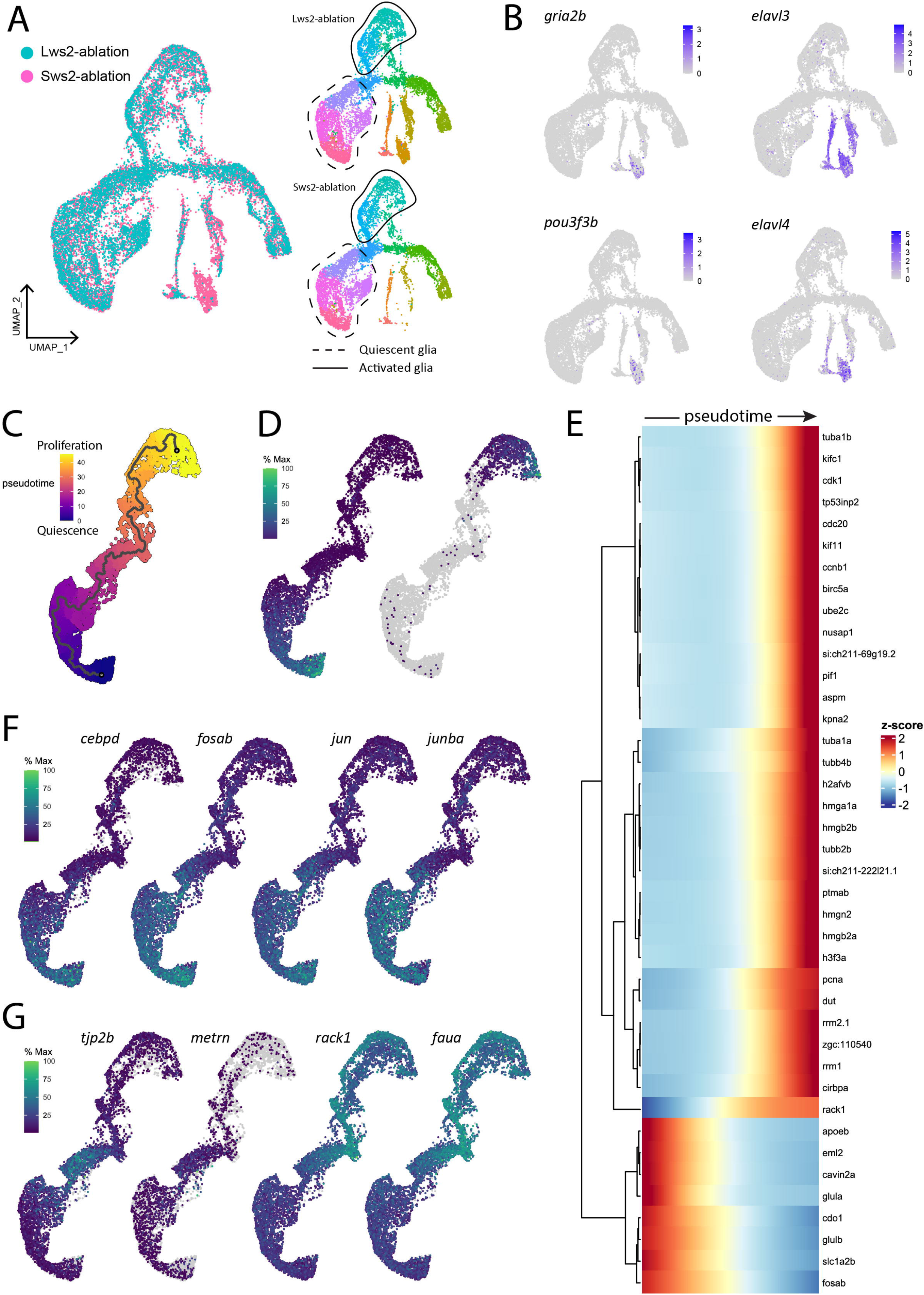
Gene expression modules in activated Müller glia following photoreceptor ablation. (A) UMAP plot of integrated Lws2 and Sws2-ablation scRNA-Seq samples. (B) Differences in clusters between photoreceptor ablation samples is associated with differentiating Müller glia-derived progenitor cells. (C) Trajectory of integrated dataset along pseudotime from quiescent (*glula*) to late proliferating (*cdk1*) Müller glia. (D) Heatmap showing the top forty differentially expressed genes along pseudotime from left to right, grouped together by gene modules. (E, F) Expression plots of differentially expressed markers along pseudotime.

Six temporally expressed gene modules along pseudotime were observed (Figure 1E). The first and second modules showed expression of genes related to mature Müller glia (*glula, glulb*, and *rlbp1a*) and retinal homeostasis (*slc1a2b, apoeb*, and *atp1a1b*) Fos and Jun-family members, transcriptional regulators that dimerise to form AP-1 transcriptional regulators involved in regulation of cell proliferation and differentiation, were also highly expressed in these Müller glia. Additionally, *cdo1* and *cebpd* associated with cell proliferation and differentiation were upregulated, as well as *mlc1a*, associated with Müller glia resting state across vertebrate species (Hoang et al., 2020). However, as Müller glia transitioned to an activated state, they repressed expression of these genes, and temporally activated a module including transient expression of genes including *notch3* and *her12, id1* (Inhibitor of DNA binding), the glial differentiation gene meteorin (*metrn*) and the tight junction protein-encoding gene *tjp2b* (Figure 1G, Supplementary Figure 1).

The third module of gene expression involved ribosomal genes *faua* and *rack1, si:dkey-151g10.6*, and Eef-family genes (*eef1g*, *eef1a1l1*, *eef1b2* and *eef2b*). These genes, which are induced as Müller glia dedifferentiate (lose glia marker expression) and enter the cell cycle, are likely important in driving this regenerative cascade (Figure 1E, G, Supplementary Figure 2). Upon entering the cell cycle, Müller glia upregulate the S-phase markers *pcna* (proliferating cell nuclear antigen) and *dut* (deoxyuridine triphosphatase), and *cirbpa* in this fourth module. Next, module five was enriched for genes relating to chromatin remodelling and the cytoskeleton, expressing histone mobility group (HMG) genes *hmgb2e, hmgn2, hmgb2b, hmga1a* as well as other histone-associated genes (*h2afvb* and *h3f3a*), alpha tubulin (*tuba1a*) and beta tubulin (*tubb2b* and *tubb4b*) genes. Finally, module six reflects Müller glia progressing into G2/M phase, with expression of *cdk1, cdc20, nusap1* and the kinesin family member genes *kifc1* and *kif11* (Figure 1E, Supplementary Figure 2). Thus, these data suggest functional competencies adopted by Müller glia that sequentially underpin gene expression as they exit their quiescent state to move towards a proliferative state.

### 3.1 Regional differences in Müller glia regenerative ability exist in the zebrafish retina

Previous studies indicate that the recruitment of Müller glia to enter this neurogenic pathway is dependent on the proximity to, and size of neural injury (Wan et al., 2014; Powell et al., 2016; Ng Chi Kei et al., 2017). However, whether all Müller glia are capable of proliferating and subsequently contributing to regeneration has not been systematically tested. We generated a variety of nitroreductase driven, metronidazole inducible ablation models that target specific photoreceptor subtypes (Figure 2A-H), which differ in their relative abundances and distribution across the larval zebrafish retina (Noel et al., 2021). These were the previously mentioned cone photoreceptor ablation models Tg(*lws2:nfsb-mCherry*) targeting long wavelength sensitive cones (Tsujimura et al., 2010) and Tg(*sws2:nfsb-mCherry*) targeting short wavelength sensitive cones (Yoshimatsu et al., 2016), as well Tg(*xops-nfsb-mCherry*) targeting rod photoreceptors (Xu et al., 2020). Additionally, we optimised a lesser injury model in Tg(*lws2:nfsb-mCherry*) fish through weaker chemical induction of photoreceptor damage that was similar in extent to the ablation observed in Sws2 and Xops ablation paradigms, allowing us to minimise differences driven by injury extent.

**Figure 2:**
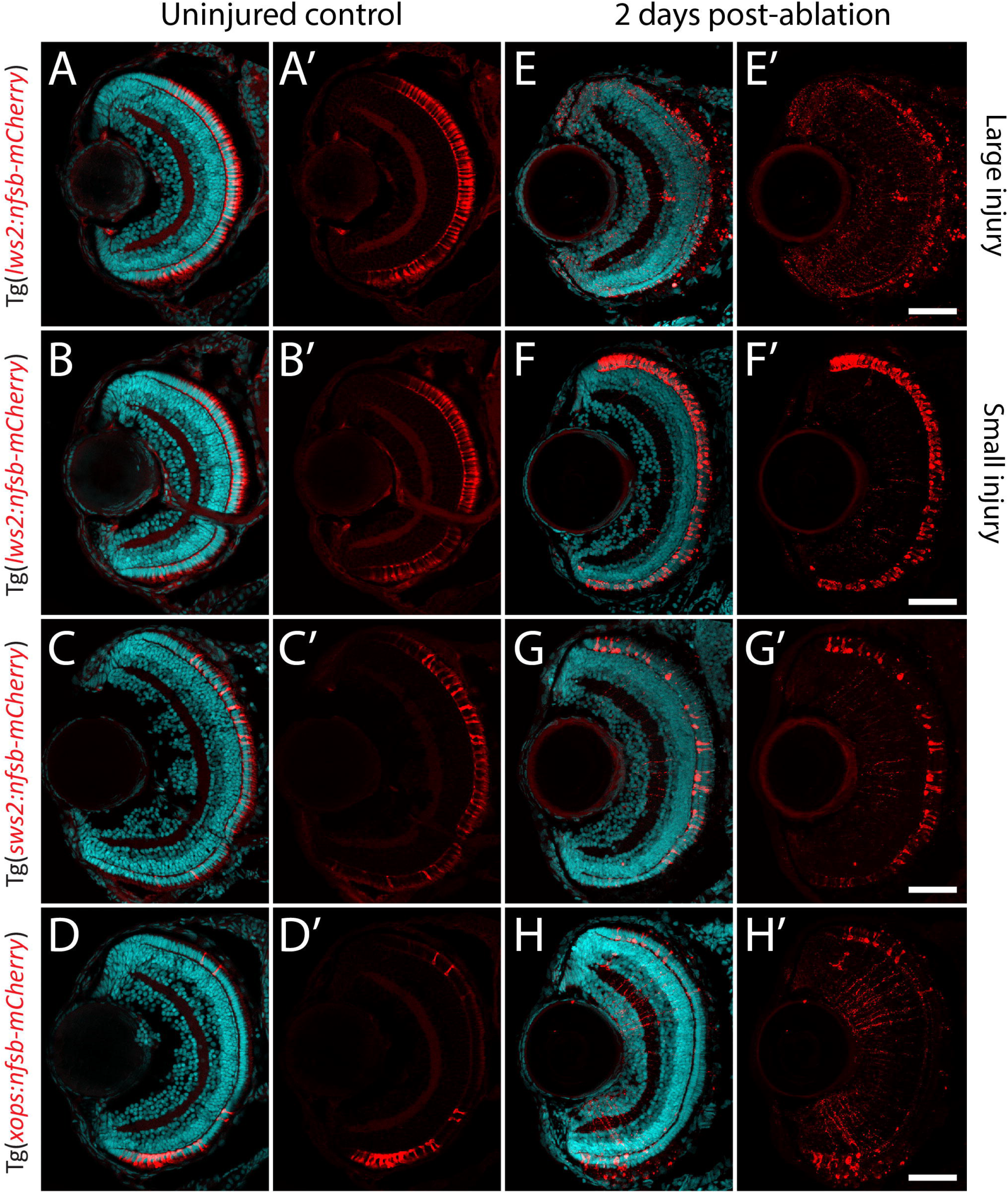
Photoreceptor ablation paradigms differing in subtype targeted, injury extent and injury location. (A-D) Distribution of mCherry positive, and nitroreductase (nfsb)-expressing long wavelength sensitive (Lws2) cones (A, B), short wavelength sensitive (Sws2) cones (C) and rod (Xops) photoreceptors (D). (E-H) Ablation of these photoreceptor subtypes is observed within 2 days exposure of metronidazole. A reduced metronidazole concentration and exposure duration leads to a smaller injury scale (B, F) compared to the original Lws2 injury (A, E). Scale bar = 50 μm.

With this approach, we quantified the percentage of Müller glia positive for proliferating cell nuclear antigen (PCNA) in our Tg(*gfap:GFP*) zebrafish crossed to either Tg(*lws2:nfsb-mCherry*), Tg(*sws2:nfsb-mCherry*) or Tg(*xops:nfsb-mCherry*) at 48 and 72 hours post-injury (hpi), as a readout of the quiescent Müller glia that were activated to drive the regenerative neurogenic response (Figure 3A-H). Following widespread Lws2-ablation (Figure 3A, B), the majority of PCNA-positive, proliferating glia were found in the central sector (24.4 ± 2.8 % at 48 hpi, 26.0 ± 3.5 % at 72 hpi) and dorsal sector (19.8 ± 2.9 % at 48 hpi, 22.1 ± 3.10 % at 72 hpi), with no significant differences between these sectors at either time point recorded. In contrast, the proportion of proliferating Müller glia in the ventral sector was significantly reduced when compared to dorsal (p < 0.001) and central (p < 0.0001) Müller glia at both 48 (2.62 ± 1.70 %) and 72 hpi (4.88 ± 2.94 %), despite Lws2 cones being abundant in this region as well. This same pattern was observed following our smaller Lws2-ablation (Figure 3C, D); dorsal (8.4 ± 2.1 %; p = 0.03) and central (12.7 ± 3.1 %; p < 0.001) Müller glia showed significantly higher proliferation than ventral Müller glia (1.2 ± 0.9 %) at 48 hpi. No significant difference in proliferation was observed at 72 hpi between Müller glia in dorsal (5.2 ± 2.2 %), central (7.5 ± 1.7 %) and ventral (1.3 ± 0.8 %) domains.

**Figure 3:**
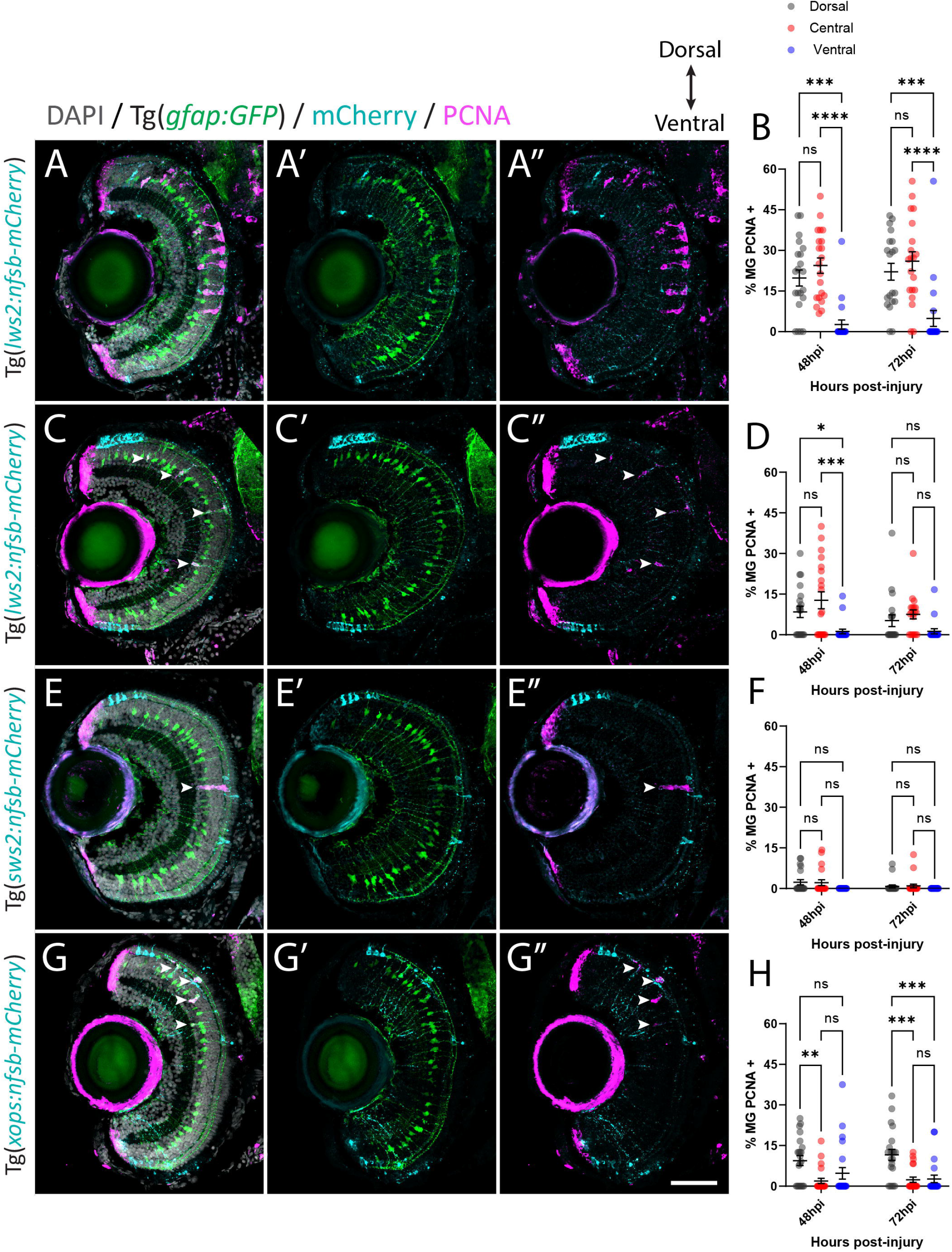
Müller glia subpopulations along the dorsal to ventral axis differ in their regenerative ability. Response of Müller glia following widespread Lws2 (A), smaller Lws2 (C), Sws2 (E) and Xops (G) ablation. Proliferation of Müller glia (labelled for PCNA; pink) expressing GFP (green), driven by the promoter for glial fibrillary acidic protein (*gfap*). Nuclei are shown in grey. All retinal sections are orientated dorsal (top) to ventral (bottom). Quantifications of PCNA-positive Müller glia for each injury at 48 and 72 hours post injury (hpi) in dorsal, central and ventral sectors (B, D, F, H). Scale bar = 50 μm.

Minimal Müller glia proliferation was observed following Sws2-ablation (Figure 3E, F), which has been previously reported in zebrafish (D’Orazi et al., 2020). PCNA-positive Müller glia were only observed at 48 hpi (2.3 ± 0.9 % and 2.2 ± 1.1 %) and 72 hpi (0.8 ± 0.5 % and 1.0 ± 0.7 %) in dorsal and central sectors of the retina respectively. Ventrally located PCNA-positive Müller glia were not observed following Sws2-ablation. After Xops ablation (Figure 3G, H), which are restricted to the ventral and dorsal regions of the retina at these larval stages, Müller glia activation occurred predominantly at the site of neural cell death consistent with previously published results (Lenkowski and Raymond, 2014). We observed only very minimal proliferation of central Müller glia (1.9 ± 1.0 % and 2.4 ± 1.0 %) at 48 and 72 hpi, respectively. Indeed, most of the proliferating Müller glia were detected in the dorsal domain (9.4 ± 1.8 % at 48 hpi and 11.6 ± 2.1 % at 72 hpi). However, ventrally located Müller glia proliferated, at a reduced capacity compared to dorsally located Müller glia (4.8 ± 2.1 % at 48 hpi; p = 0.1 and 2.7 ± 1.4 % at 72 hpi; p < 0.001).

### 3.2 Regardless of photoreceptor subtype ablated, the majority of Müller glia contribute to debris phagocytosis

Following rod ablation, proliferative Müller glia were predominantly located in the peripheral (dorsal and ventral) retina, which matched the overall distribution of rod photoreceptors (mCherry-positive) at this age (Figure 2D, H). Thus, we assessed if the differences in capacity for activation of Müller glia are influenced by signals local to the site of injury. Specifically, we investigated phagocytosis of dying cells and their debris, which can influence behaviour Müller glia following injury (Sakami et al., 2019; Lew et al., 2022) including Müller glia proliferation (Bailey et al., 2010; Nomura-Komoike et al., 2020). For this, we quantified the number of mCherry containing presumed phagocytic Müller glia and compared this to the Müller glia re-entering the cell cycle (PCNA-positive) after widespread Lws2, smaller Lws2 and rod ablation, as these resulted in a substantial Müller glia proliferative response (Figure 4). As shown, we found that more Müller glia contained mCherry-positive debris than expressed PCNA in the small Lws2 cone and rod ablation studies, yet more Müller glia re-entered the cell cycle in the widespread Lws2 cone ablation (Figure 4A – L). In all cases, we found a significantly higher proportion of phagocytic (mCherry-positive) Müller glia within the proliferative population compared to the total Müller glia cohort at 48 hpi (22 % vs 13 % Lws2 big injury; 69 % vs 48 % Lws2 small injury; 87 % vs 46 % Xops injury; p < 0.0001) and 72hpi (25 % vs 14 % Lws2 big injury; 62 % vs 33 % Lws2 small injury; 76 % vs 42 % Xops injury; p < 0.0001)(Figure 4H, J, L). Thus, Müller glia that phagocytose substantial amounts of dying photoreceptors are significantly more likely to respond to injury and mount a regenerative response, consistent with phagocytosis being one of the cellular processes that may convey injury signals to recruit Müller glia for regeneration.

**Figure 4:**
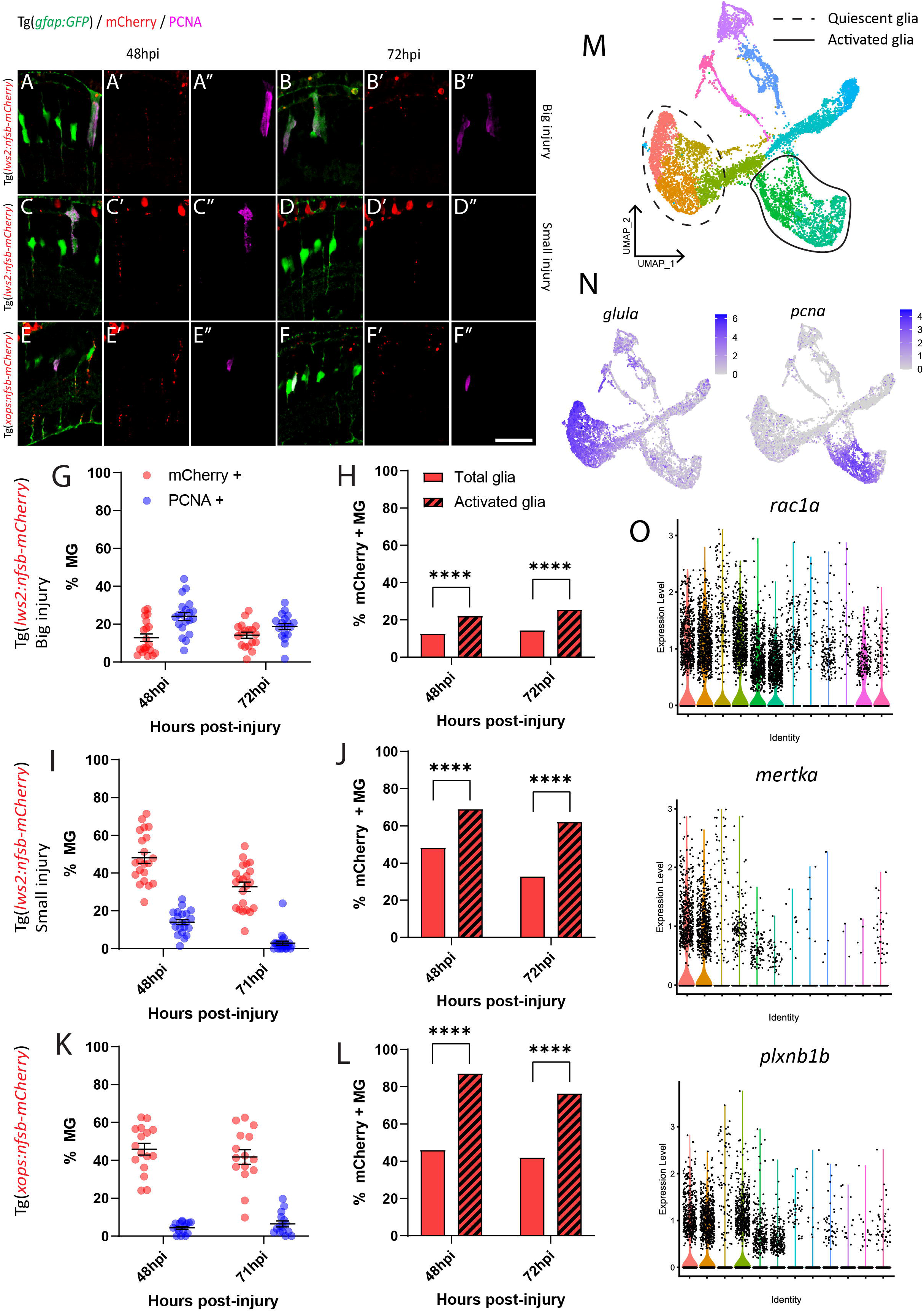
Investigation of phagocytosis and proliferation by Müller glia following photoreceptor ablation. (A-F) Micrograph images of Müller glia (Gfap-positive; green) detecting proliferative marker PCNA (pink) and photoreceptor debris (red) across three photoreceptor ablation paradigms. (G, I, K) Quantifications of the percentage of debris-containing, and proliferating Müller glia. (H, J, L) Cell counts presented as proportions of debris-containing Müller glia in PCNA+ and PCNA-populations. (M) UMAP plot of quiescent (dotted line) and proliferative (solid line) Müller glia clusters expressing *glula* and *pcna* respectively (N) from Lws2 ablation scRNAseq sample. (O) Violin Plots of the expression of phagocytosis associated markers *rac1a, mertka* and *plxnb1b*. * = p ≤ 0.05, ** = p ≤ 0.01, *** = p ≤ 0.001, **** = p ≤ 0.0001.

Additionally, we wanted to detect even minute levels of mCherry debris, which may not have been visible after standard immunofluorescence (and antigen retrieval) methodology. This was conducted through repeating analysis of mCherry-positive and PCNA-positive Müller glia using an mCherry antibody with all injury paradigms (Supplementary Figure 3). Surprisingly, we indeed found that the majority of Müller glia contained mCherry-positive debris when stained with antibodies against mCherry at 24 hpi (99.4± 0.4 %, 80.7 ± 7.6 %, 73.2 ± 11.2 % and 74.8 ± 6.2 %), 48 hpi (99.7 ± 0.3 %, 88.9 ± 5.9 %, 85.1 ± 5.3 % and 81.3 ± 3.7 %), 72 hpi (99.6 ± 0.4 %, 85.5 ± 4.3 %, 81.5 ± 5.5 % and 81.2 ± 4.4 %) and 96 hpi (95.2 ± 2.8 %, 73.4 ± 5.0 %, 78.8 ± 5.8 % and 73.4 ± 4.1 %) in our widespread Lws2, smaller Lws2, Sws2 and Xops-ablations respectively (Supplementary Figure 3A-H). Thus, the proportion of proliferating (activated) and total (including quiescent) Müller glia that contained any mCherry debris was both very high. Thus, while we did still observe that proliferating Müller glia were significantly more likely to contain mCherry debris in our smaller Lws2 ablation at 72 hpi (p = 0.04), and rod photoreceptor ablation models at 48 hpi (p = 0.03, Supplementary Figure 3I – K), phagocytosis does not seem to be sufficient for Müller glia activation. Correlating this to our scRNAseq data, we found that phagocytosis-associated markers are also upregulated following injury specifically in both quiescent and proliferating Müller glia cell clusters (Figure 4M – O), consistent with this cellular process preceding de-differentiation and cell cycle re-entry. Thus, phagocytosis alone does not seem to be sufficient as a functional process to activate the injury response in glia, but is significantly correlated to Müller glia activation.

### 3.3 Mature zebrafish Müller glia show heterogeneous gene expression in the absence of neural injury

Given that we saw spatially distinct activation of Müller glia that could not be explained by site of cell ablation or phagocytic activity alone, we assessed for any heterogeneity in gene expression patterns prior to injury in the mature, quiescent Müller glia population. To achieve this, we performed scRNA-seq of FACS-Müller glia from 6 days post fertilization (dpf) Tg(*gfap:GFP*) zebrafish larvae. UMAP construction and unsupervised clustering revealed 11 populations of Müller glia and Müller glia-derived cells. Of these, 6 neighbouring clusters were very similar (Figure 5A) and showed consistent expression of common genes typical of quiescent Müller glia, such as high expression of glial markers *glula, gfap* and *rlbp1a* (Figure 5B). The remaining 5 clusters did not represent the quiescent populations we were focusing on (marked in grey in Figure 1A). These cell clusters expressed markers of proliferation (*pcna*) or neural differentiation (*crx*, *neurod4, elavl3, elavl4*) or intermediate states (*her4.1, igfbp5b*) (Figure 5C). These clusters likely encompass either young Müller glia exiting a progenitor state or a subset of Müller glia that sporadically enter the cell cycle to generate rod photoreceptors (Bernardos et al., 2007; Ng et al., 2014). Additionally, one cluster displayed high expression of genes enriched for terms suggestive of stress response (Supplementary 4A), such as *hsp90aa1.2, hsp70.3 and hsd11b2* (data not shown). From these results, we excluded these clusters for further analysis, and we focused on clusters representing mature quiescent glia (C1-C6). We performed hierarchical clustering analysis of top-ranking expressed genes clusters C1 to C6 and identified key similarities (reflected in their close proximity on the UMAP plot), as well as subtle, but robust differences between these cell clusters (Figure 5A; Supplementary Figure 4B). We found an unexpected diversity of quiescent Müller glia gene expression states within the uninjured retina (Figure 5D, E). While these six clusters were overall grouped together due to their overwhelming similarities as quiescent glia, robust differences were observed in the gene lists that led to unbiased sub clustering.

**Figure 5:**
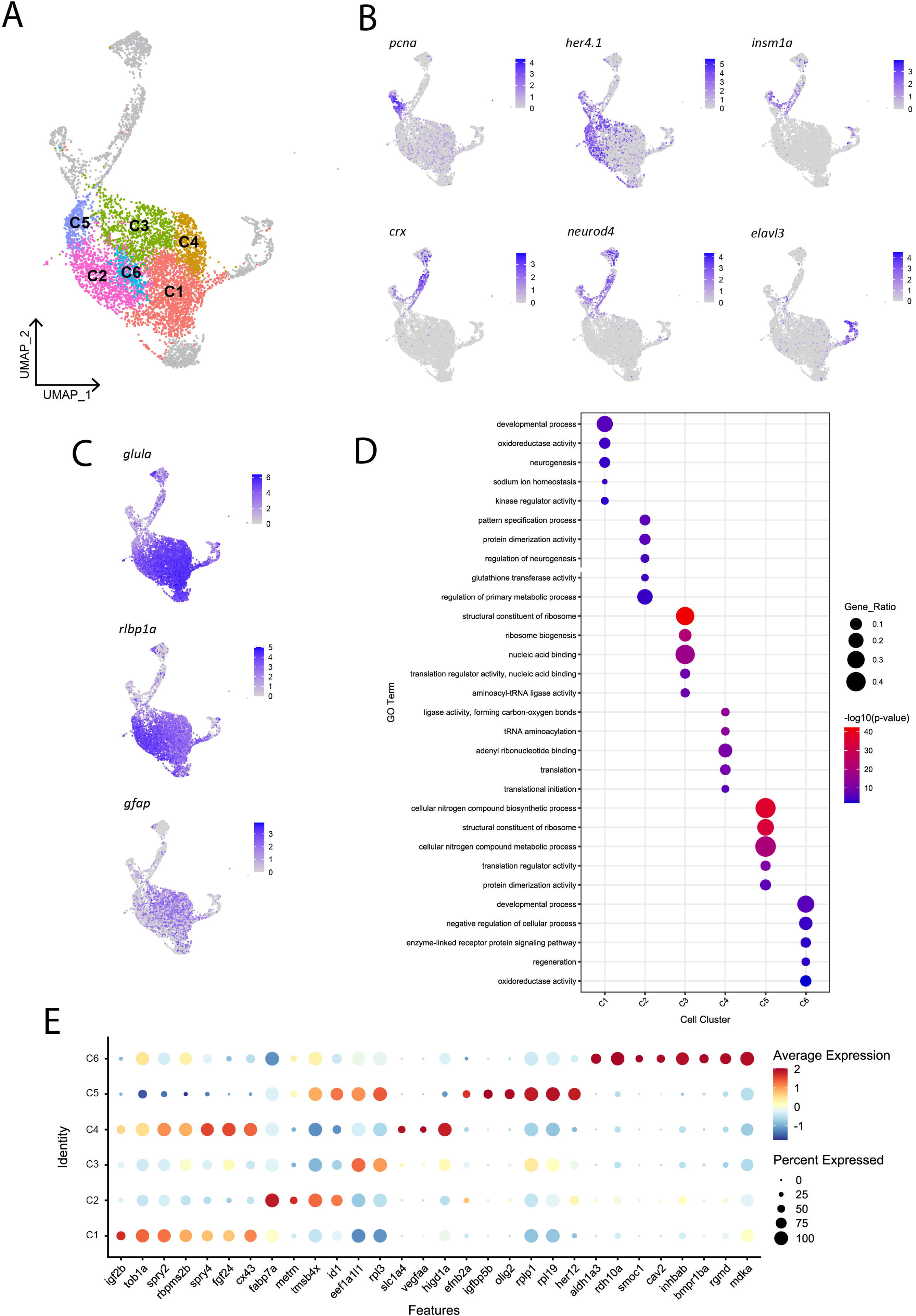
Heterogeneity exists in Müller glia of the uninjured zebrafish retina. (A) UMAP of FACS Müller glia and Müller glia-derived cells from the uninjured zebrafish retina at 9 days postfertilisation (dpf), revealing six clusters of quiescent Müller glia (C1-C6). B) Expression plots of proliferating (*pcna*), immature cells (*her4.1* and *igfbp5b*), and differentiating neurons (*crx, neurod4* and *elavl3*). (C) Expression of mature Müller glia markers in the identified quiescent Müller glia. (D) Enrichment term analysis summarized as 5 gene ontology (GO) terms for each clusters C1-C6. Circle size depicts the gene ratio and circle colour depicts the adjusted negative log *p*-value of significance. (E) Expression of top genes across clusters C1-C6. Circle size and circle colour represents percentage of expressing cells per cluster and average log-fold expression value respectively.

Cluster C1 displays enrichment for terms including developmental process, neurogenesis and kinase regulator activity, in addition to retinal homeostasis (oxidoreductase activity and sodium ion homeostasis). This cluster expresses high levels of the *igf2b* (insulin-like growth factor 2b). Among other highly expressed genes in this cluster includes *tob1a* involved in development of dorsal structures (Xiong et al., 2006), fibroblast growth factor 24 *fgf24*), the negative regulators of receptor tyrosine kinase signalling *spry2* and *spry4* (Felfly and Klein, 2013), the RNA-binding protein *rbpms2b*, and gap junction protein *cx43*. Unique to cluster C2 is expression of the glial differentiation gene meteorin (*metrn*) and fatty acid binding protein 7a (*fabp7a*). Cluster C2 also expresses the ID signalling gene *id1* and thymosin beta-4 (*tmsb4x*) and enrichment for terms relating to pattern specification process and regulation of neurogenesis. Clusters C3, C4 and C5 were enriched for terms relating to ribosome activity, including ribosome biogenesis, translation initiation and structural constituent of ribosome. Cluster C3 expressed genes *eef1a1l1* (translation elongation factor) and *rpl3* (60S ribosomal protein L3). Cluster C4 expressed growth factor signalling-associated genes *fgf24* and *spry4* that were shared with cluster C1, as well as unique genes encoding the amino acid transporter *slc1a4*, vascular endothelial growth factor Aa (*vegfaa*) and hypoxia inducible domain family member 1a (*higd1a*). Additionally, cluster C5 is enriched for terms related synthesis of biological compounds and metabolism, and cells in this cluster express the bHLH transcription factor *olig2, igfbp5b* (insulin-like growth factor binding protein 5b), *her12* and ephrin receptor-binding ligand *efnb2a*. Despite containing a relatively smaller proportion of cells, the central cluster C6 is distinct in its gene expression profile, displaying enrichment for terms relating to development, regeneration and cellular signalling, expressing genes of the retinoic acid pathway (*aldh1a3, rdh10a*), and TGFβ superfamily (*inhbab, bmpr1ba, rgmd*), as well as the caveolar-associated gene *cav2*, basement membrane protein-encoding gene *smoc1* and midkine-a (*mdka*). In summary, we have identified the following putative quiescent Müller glia subpopulations: Two subpopulations associated with fibroblast growth factor (Fgf) signalling, which can be distinguished based on the presence (C1) or absence (C4) of *igf2b*, a *fabp7a-*expressing subtype (C2), Müller glia strongly associated with protein production (C3 and C5) and finally a Müller glia subtype associated with retinoic acid signalling (C6). Whether these clusters represent stable specialisations in the terminal differentiation of this cell population to support their diverse roles in maintaining retinal homeostasis, fluid states through which individual glia stochastically transition through, or are indicative of varying developmental ages of Müller glia remains an interesting avenue to explore.

### 3.4 Molecularly distinct quiescent Müller glia subpopulations show a distinct spatial distribution

Without spatial or temporal information on individual Müller glia, the scRNA-seq data cannot distinguish whether these differing clusters represent distinct Müller glia subpopulations. If the molecular heterogeneity was linked to specific functions or represent stochastic fluid transition states, we hypothesized to find glia of different gene expression clusters distributed either uniformly or stochastically across all retinal regions. In contrast, we postulated that if the clustering was dependent on the age of Müller glia, these clusters would manifest as a central-to-peripheral distribution pattern in the retina. The retina of fish and amphibians exhibits indeterminate eye growth, with new neurons and glia continuously added from the peripheral ciliary margin zone (CMZ) into the growing central retina (Centanin et al., 2011; Fischer et al., 2013; Centanin et al., 2014). Therefore, older Müller glia are found more centrally than those in peripheral regions. With these possibilities in mind, we assessed the distribution of the distinct clusters both in the developing and adult retina. We performed RNAscope *in situ* hybridisation on candidate genes *efnb2a* (clusters C2 and C5), *fgf24* (clusters C1 and C4), and *rdh10a* (cluster C6). Unexpectedly, we found the presence of these markers in regionally distinct Müller glia within the dorsal (21%), central (50%) and ventral retina (19%) respectively (Figure 6A-D). As expected, PCNA-positive cells were found in the peripheral ciliary margin zone, likely from which *igfbp5b*-positive, *her4.1*-positive immature Müller glia are derived from (Figure 6D). Furthermore, this distribution was maintained throughout maturation, with a similar pattern observed at 12 months post-fertilisation (Supplementary Figure 5A). Thus, our clustering analysis has led to the identification of genes that mark regional populations of Müller glia in a pattern that persists with ageing.

**Figure 6:**
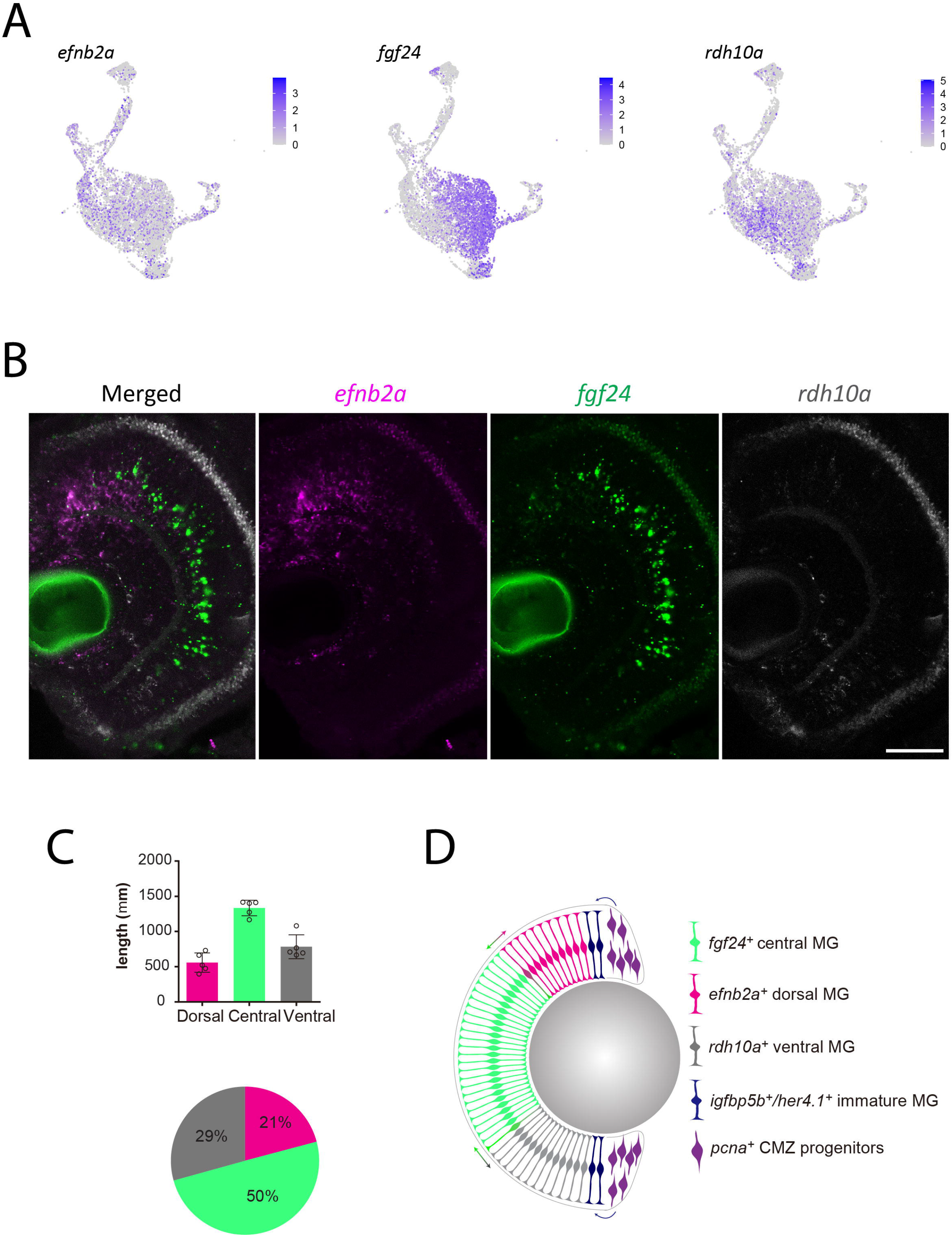
Molecularly distinct Müller glia subpopulations differ in their spatial location. (A) Expression plots of key markers *efnb2a, fgf24* and *rdh10a*. B) RNAscope *in situ* hybridization of genes presented in (A), highlighting their location throughout the dorsal (top) to ventral (bottom) retina. Scale bar = 50 μm. (C) Length (μm) in the retina of 12 month-old zebrafish retina domains of *in situ* labelling. (D) Schematic summary of the location of *efnb2a* (pink), *fgf24* (green), *rdh10a* (grey)-expressing Müller glia, in addition to *igfbp5b/her4.1*-expressing immature Müller glia and *pcna*-expressing ciliary margin zone progenitor cells.

We next assessed the pattern of Müller glia recruitment following Lws2 cone photoreceptor ablation using Tg(*lws2:nfsb-mCherry*) in adulthood. We first characterised the distribution of Lws2 cones and Müller glia in the uninjured adult fish (Figure 7A). Consistent with our data on larvae, both Lws2 cones and Müller glia are evenly distributed, with some additional cells in the central retina (Figure 7B). Following ablation, we used *fgf24* and *rdh10a* as regional markers of spatially segregated Müller glia subpopulations. Consistent with the larval retina (Figure 3), we saw a specific bias of Müller glia in the dorsal and central *fgf24+* glia regions, with very little activation and cell cycle entry activity (marked by PCNA immunostaining) in the ventral *rdh10a+* regions (Figure 7C). Thus, even in the context of widespread photoreceptor injury across the retina, Müller glia show region-specific heterogeneity in their capacity to proliferate, and this heterogeneity persists from larval to adult stages. Thus, our clustering analysis has led to the identification of genes that mark regional populations of Müller glia with differing probability to contribute towards the regenerative response.

**Figure 7:**
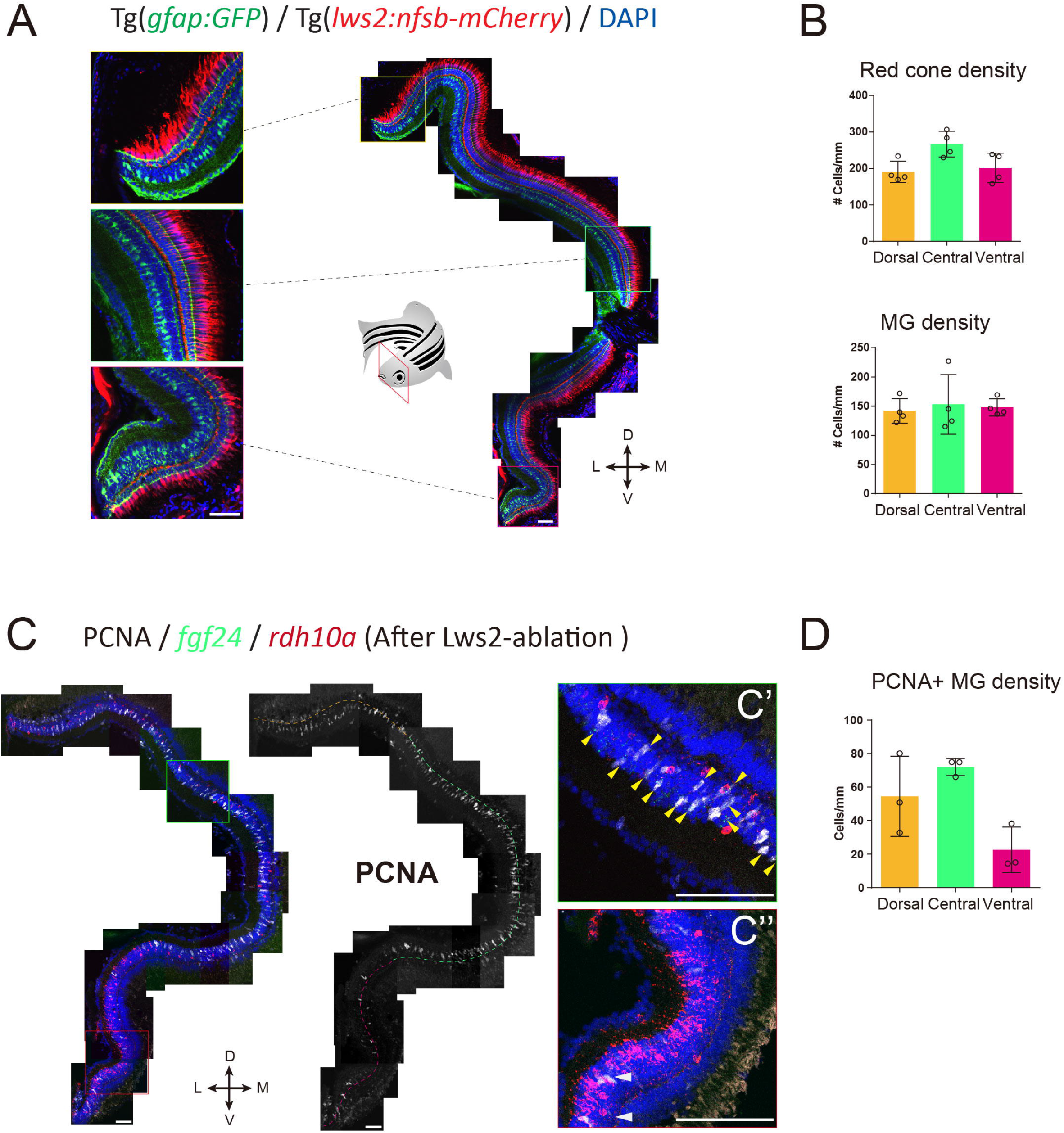
*fgf24+* Müller glia are primed to proliferate upon red cone ablation in adult zebrafish. (A) The distribution of red (Lws2) cones and Müller glia in the adult retina. (B) Quantifications of the density of red cones and Müller glia in each spatial domain. (C) Red cone ablation significantly induces central Müller glia to proliferate and label for PCNA. Yellow arrows indicate *fgf24+/PCNA+* Müller glia (C’), while white arrows indicate *rdh10a+/PCNA+* Müller glia (C”). (D) Quantifications of the density of PCNA+ Müller glia in each spatial domain. Scale bar = 50 μm.

### 3.5 Distinct recruitment of Müller glia subpopulations following neural ablation

We found that Clusters C1 to C6 featured GO terms for mature Müller glia functions including neuroprotection, metabolism and ion homeostasis (Newman et al., 1984; Resta et al., 2007; Bringmann and Wiedemann, 2012; Furuya et al., 2012; Reichenbach and Bringmann, 2016). Yet, wanted to determine whether these populations differed in their responsiveness to injury or regenerative potential. Cluster C6 revealed many markers that are known to influence stem cell renewal (Figure 5E), including the bone morphogenetic protein receptor *bmpr1ba*. (Bond et al., 2012; Wang et al., 2014), TGF-β family ligands *inhbab* and *rgmd* (Heldin et al., 2009), retinoic acid pathway proteins *aldh1a3* and *rdh10a* (Blum and Begemann, 2012), and the cytokine *mdka* (Ang et al., 2020). Immunofluorescent labelling of Bmpr1ba/b showed localization in neurons (DAPI-positive, Gfap-negative cells) and Müller glia (DAPI-positive, Gfap-positive cells) in Tg(*gfap:GFP*) zebrafish (Supplementary Figure 6). We observed labelling in Müller glia in the ventral retina, as well as in the central sector, with minimal labelling present on Müller glia in the dorsal retina. This supports the association of cluster C6 with glia in the ventral domain of the retina. Furthermore, the restriction of these proliferation-associated genes to a subset of Müller glia motivated us to explore potential regional differences in Müller glia proliferation.

To explore this regional heterogeneity in Müller glia activation, we conducted integration and re-clustering analysis of cells collected from our no-ablation and Lws2-ablation scRNA-seq datasets (Figure 8A). There, we identified 4 main quiescent Müller glia populations marked by *fgf24*, *efnb2a* and *rdh10a* expression as genes most uniquely expressed for these quiescent Müller glia clusters, consistent with observations from the uninjured sample alone (Figure 8B, C). Notably, *fgf24*-positive Müller glia spanned two clusters distinguished by the expression of the insulin growth factor-encoding gene *igf2b* and the transmembrane glycoprotein-encoding gene *cd99*. Curiously, *cd99* exhibited greater expression following photoreceptor ablation. Furthermore, levels for immune-associated genes also changed following photoreceptor ablation, including the chemokine genes *cxcl18b* and *cxcl14*, which were higher in expression following photoreceptor ablation when compared to the uninjured dataset in these quiescent Müller glia clusters (Supplementary Figure 7A), specifically in this grouping of *cd99*-positive cells (Supplementary Figure 7B). As such, certain Müller glia may be functionally specialized to signal to immune cells.

**Figure 8:**
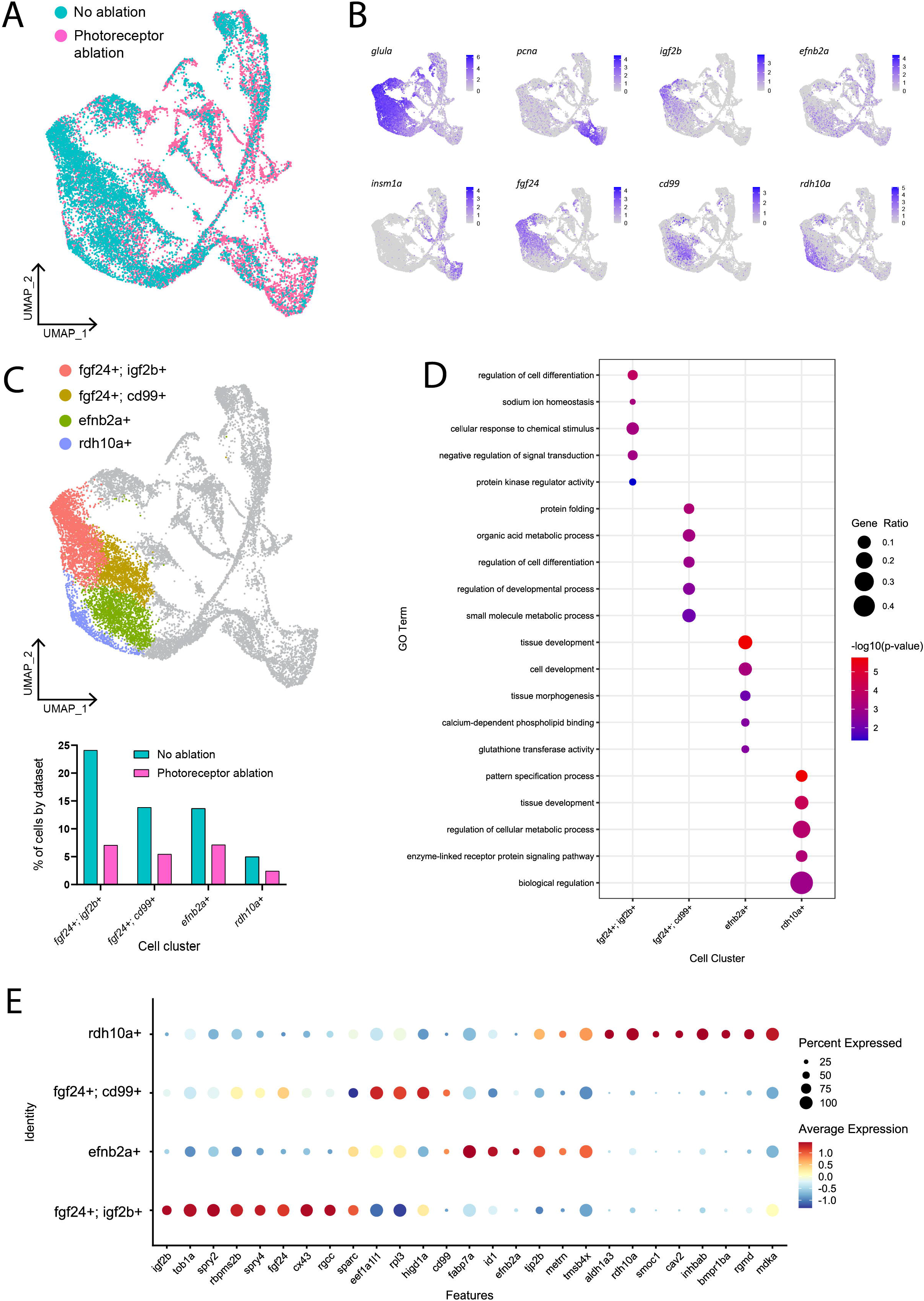
Müller glia heterogeneity persists in the presence of photoreceptor ablation. (A) UMAP plot of FACS Müller glia with integration of uninjured (no ablation) and photoreceptor ablation single cell RNA-sequencing samples. (B) Expression plots of quiescent (*glula*), proliferating (*pcna*) and differentiating (*insm1a*) cells, as well as key markers distinguishing quiescent Müller glia clusters. (C) UMAP plot highlighting quiescent Müller glia clusters that can be distinguished based on the expression of the following markers: *fgf24, igf2b, cd99, efnb2a* and *rdh10a*. Graph indicates the percentage of cells belonging to each cell cluster and the proportion between each sample. (D) Enrichment term analysis of genes expressed in each quiescent cluster and the gene ontology (GO) terms relating to these genes. Circle size and circle colour depict gene ratio and negative log-adjusted *p*-value respectively. (E) Expression of key markers linked to quiescent Müller glia heterogeneity that persist in the presence of photoreceptor ablation. Circle size indicates percentage of cells expressing the relevant gene and circle colour indicates the average log-fold expression value.

The range of molecular profiles of quiescent Müller glia remained following neural injury. Gene ontology analysis revealed that the differentially expressed genes in *fgf24; igf2b-*positive, *fgf24; cd99*-positive*, efnb2a-*positive, and *rdh10a*-positive clusters were enriched for biological processes relating to development, regeneration, or differentiation (Figure 8D, E). As the additional branches of activated Müller glia and differentiating progenitors in our photoreceptor ablation sample must come from the original population of quiescent Müller glia from the uninjured sample, we compared whether all the quiescent Müller glia populations were recruited equally. The greatest differences observed in the proportion of Müller glia between the injury and non-injured control was in the following order: *fgf24; igf2b-positive* (70.7%), *fgf24; cd99*-positive (60.6%) clusters, followed by similar differences in *rdh10a*-positive (51.6%) and *efnb2a*-positive (47.9%) clusters (Figure 8C). Thus, within the *fgf24-*positive population of glia, proliferative ability may be influenced by expression of *igf2b*.

## 4 Discussion

In this study we have identified an unexpected heterogeneity in mature quiescent Müller glia of larval zebrafish that is marked by differential gene expression, and which persists over age. Some of this heterogeneity is linked to specific homeostatic glia functions; the expression of *vegfaa* and *higd1a* share protective roles within both Müller glia and surrounding retinal neurons, regulating vascular function in both normal and hypoxic conditions (*vegfaa*) and protecting from reactive oxygen species during metabolic stress (*higd1a*) (Hayashi et al., 2015; Le, 2017). Their expression in a subset of Müller glia may indicate functionally specialised Müller glia subpopulations. Additionally, our analyses revealed six distinct clusters that differed in their expression of genes primarily relating to protein synthesis, development and tissue patterning. This heterogeneity was maintained into adult ages, and persisted in the presence of injury, with clear distinction in expression of the markers *efnb2a*, *fgf24* and *rdh10a*, which all play roles in dorsoventral patterning in the retina (Sen et al., 2005; Sakuta et al., 2006; Atkinson-Leadbeater et al., 2014; Zhang et al., 2016). These markers identified Müller glia subpopulations along the dorsal ventral axis, whereby Müller glia that re-entered the cell cycle were primarily restricted to central (expressing *fgf24*) and dorsal (expressing *efnb2a*) domains of the retina when challenged with a variety of neural ablation paradigms that model photoreceptor degeneration (Baumgartner, 2000; Kalloniatis and Fletcher, 2004; Hamel, 2006; Lamba et al., 2008; Ferrari et al., 2011; Masland, 2012; Xu et al., 2020; Fleckenstein et al., 2021; Sarkar et al., 2022). Our results highlight the importance of the intrinsic molecular state in dictating whether Müller glia will respond to injury and subsequently be recruited to contribute to regeneration.

Due to their typical arrangement, morphology, and conserved expression of markers, the regional heterogeneity we identified in Müller glia was surprising. As such, our study offers clues into the molecular states for spatially restricted, quiescent Müller glia that drive cellular reprogramming, cell cycle entry and neuronal regeneration. Indeed, when comparing uninjured with photoreceptor-ablated retinas, resting Müller glia expressing both *fgf24* and *igf2b* were preferentially recruited in the regenerative response. Both insulin (IGF) and fibroblast (FGF) growth factors are necessary for stimulating Müller glia proliferation in zebrafish (Wan et al., 2014), chick (Fischer et al., 2002), and mouse (Karl et al., 2008) retina. Within the highly regenerative zebrafish, multiple Fgf members have been shown to play distinct roles in neural regeneration (Goldshmit et al., 2018). Importantly, in both zebrafish (Wan and Goldman, 2017) and mammals (Close et al., 2006), the age of the retinal tissue can impact on the effect that growth factors have on Müller glia proliferation. Our data revealed that the negative downstream feedback regulators of FGF signalling *spry2* and *spry4* (Christofori, 2003), which are involved in vertebrate eye development (Taniguchi et al., 2007; Kuracha et al., 2011) are also associated with *fgf24-expressing* Müller glia. SPRY2 and SPRY4 in humans are expressed in human embryonic stem cells (hESCs), with SPRY2 documented as having both pro (Felfly and Klein, 2013) and anti-proliferative (Zhang et al., 2005) effects. Thus, this prompts investigation of these insulin and fibroblast growth factor signalling members in the Müller glia proliferative response.

In our analyses, ventrally located, *rdh10a*-expressing Müller glia undergo minimal proliferation, even when cell death is localized to the ventral domain. We detected a network of genes which we interpret to be relevant to maintaining Müller glia quiescence, however it remains unclear as to whether these genes are permissive or restrictive to Müller glia activation in the presence of injury. Multiple members of the TGFβ signalling pathway were strongly expressed in the *rdh10a-*positive Müller glia population, including the bone morphogenetic protein receptor *bmpr1ba*, and activin ligands *rgmd*, *rgmb* and *inhbab*. This pathway plays diverse roles through activation of intracellular signalling molecules called Smads, which range from cell proliferation to growth arrest, depending on ligand-receptor interactions, the target cells involved and the tissue environment (Itoh and ten Dijke, 2007; Corradini et al., 2009; Wu et al., 2012). In the retina, Tgfβ target genes show an immediate upregulation following retinal injury. However, Tgfβ signalling can have differing effects on Müller glia depending on the timing of expression; Tgfβ signalling is necessary for the expression of pro-regenerative genes immediately after injury, yet inhibition of Tgfβ during the regenerative phase leads to increased proliferation (Lenkowski et al., 2013; Tappeiner et al., 2016; Lee et al., 2020b; Sharma et al., 2020). Interestingly, the activin ligand-encoding gene inhibin subunit beta (INHB) is associated with quiescent states of chick and zebrafish Müller glia (Hoang et al., 2020), but an activated or gliotic state in mammalian Müller glia (Hoang et al., 2020; Conedera et al., 2021). One method of regulating this pathway is through endocytosis of transmembrane receptors through invaginated plasma membrane pits called caveolae, with associated proteins including *cav2* and *cav1*, which were unique to this cluster (Anderson et al., 1992; Di Guglielmo et al., 2003; Hartung et al., 2006). While these studies have investigated Müller glia quiescence and activation in the total Müller glia population, our study is the first to investigate differences within subsets of these Müller glia.

The relevance of heterogeneous molecular states and injury responsiveness of mature Müller glia subpopulations in retinal homeostasis and vision remains unclear. We found that Müller glia age was not the key determinant of the different clustering. There may be a functional requirement of maintaining retinal circuitry (through neural regeneration) of this central/dorsal region of the retina, as loss of photoreceptors in these regions may have greater consequences to the visual field than in the ventral retina. We identified distinct domains of *fgf24* and *rdh10a-*expressing Müller glia in the zebrafish retina. Controlled expression of Retinoic acid and FGF signalling factors in the chick retina defines a region of high visual acuity (da Silva and Cepko, 2017). In our analysis, the definition of the ventral, *rdh10a*-associated region likely involves a variety of patterning genes such as the transducer of ErbB2 gene *tob1a*, unique to *fgf24*-expressing Müller glia in our dataset, which is known to inhibit Smad-mediated TGF-β signalling (Xiong et al., 2006). Whether prioritsing neuron replacement in distinct retinal regions can influence recovery of vision in the zebrafish is an area that remains unexplored. Despite retinoic acid signalling associated with improved regeneration in a variety of vertebrate species (Duprey-Díaz et al., 2016; Todd et al., 2018; De La Rosa-Reyes et al., 2021), we show that its expression in quiescent Müller glia is associated with a reduced likelihood of activation. Therefore, functional manipulation of these candidate markers may reveal intrinsic barriers to Müller glia activation. Whether these same molecular states govern mammalian Müller glia quiescence is an important avenue to investigate.

In our integrated dataset, *fgf24*-expressing Müller glia were divided into two cell clusters based on the expression of the cell surface glycoprotein-encoding gene *cd99*, which is involved in leukocyte migration and endothelial cell adhesion (Schenkel et al., 2002; Tanaka, 2016), as well as inhibiting tumour growth through suppressing EGFR signalling (Lee et al., 2020a). We observed the expression of chemokines *cxcl18b* and *cxcl14* in this grouping of *cd99-*positive cells, which all exhibited increased expression levels in Müller glia from the photoreceptor ablation dataset. The inflammatory chemokine Cxcl18b is a neutrophil-specific chemoattractant in zebrafish (Torraca et al., 2017;Sommer et al., 2020) and *cxcl18b* is upregulated following ablation of the RPE (Leach et al., 2021). CXCL14 is expressed by both immune and non-immune cells in humans and is important for leukocyte migration and differentiation (Lu et al., 2016). In zebrafish, upregulation of *cxcl14* has been observed in reactive oligodendrocyte progenitor cells following spinal cord injury (Tsata et al., 2020). The expression of immune-associated markers by Müller glia was intriguing as the presence of immune cells can influence Müller glia proliferation (Bradley, 2008; Nelson et al., 2013; Conner et al., 2014; Fischer et al., 2014; Gallina et al., 2014; White et al., 2017; Conedera et al., 2019; Mitchell et al., 2019; Todd et al., 2020; Leach et al., 2021). While we also observed a recruitment of L-plastin labelled leukocytes to the photoreceptor layer following ablation, in the absence of Müller glia, this migration to the photoreceptor layer remained unchanged (data not shown). Furthermore, contrary to what is known, elimination of mpeg1-positive macrophages and microglia did not enhance the Müller glia proliferative response following lws2-photoreceptor ablation (data not shown). Whether this is due to differences in the specific photoreceptor subtype eliminated, or the developmental age at the timing of ablation requires further investigation. Functional manipulation of these immune signalling genes following retinal injury will reveal their ability to facilitate immune cell-Müller glia communication and whether this can improve Müller glia proliferation.

In this study we showed that regardless of the injury extent, Müller glia effectively phagocytosed photoreceptor debris, and expression of phagocytosis markers *rac1a, mertka* and *plxnb1b* were detected in our scRNAseq samples. However, this cellular engulfment, whilst correlated, did not seem sufficient to cause Müller glia activation. Müller glia are able to sense injury cues released by dying neurons, with detection of iron, ADP and TNFalpha implicated in Müller glia proliferation (Nelson et al., 2013; Conner et al., 2014; Medrano et al., 2017; Iribarne et al., 2019; Boyd and Hyde, 2022). Therefore, whether the intrinsic molecular state of Müller glia can influence their receptiveness to these “pro-regenerative”signals presents an area of future research. Despite Müller glia proliferation occuring in the dorsal and central retina following sws2 photoreceptor ablation, this proliferation was minimal, which is consistent with recently published data (D’Orazi et al., 2020). Müller glia can sense changes to neural transmission, whereby mimicking an absence of neurotransmission through inhibition of GABA signalling can stimulate Müller glia proliferation (Rao et al., 2017;Kent et al., 2021). This inability to respond to loss of Sws2 photoreceptors may be attributed to these cells not being functionally mature at the timing of injury from 4 – 6 dpf. However, through single cell analysis, the gene expression modules involved in Müller glia activation were shared with that following Lws2 ablation, which induces a greater Müller glia proliferative response.

Through trajectory analysis we identified three main states that Müller glia achieve to proliferate. Intially, we see an upregulation of Jun and Fos-encoding genes, which form AP-1 transcriptional complexes that control cellular processes including proliferation. Evidence of stimulation (c-Jun) and repression (JunB) of cell cycle-related gene expression has been observed (Schreiber et al., 1999; Shaulian and Karin, 2001; Zenz et al., 2008), with ectopic expression of these AP-1-related genes actually leading to regeneration defects in the axolotl (Sabin et al., 2019). Furthermore, AP-1-related genes are upregulated immediately post neural injury (Hui et al., 2014; Hoang et al., 2020), suggesting an initial requirement of these genes in Müller glia proliferation. Expression of *cebpd* was also upregulated at this initial phase. *cebpd* (CCAAT enhancer binding protein delta) while being associated with a reactive phenotype of mammalian astrocytes, Müller glia and immune cells (Roesch et al., 2012; Spek et al., 2021)(Wang et al., 2016), is also involved in proliferation (Ko et al., 2015) and may be crucial in the zebrafish Müller glia proliferative response.

A transient reactive phase has been observed in adult zebrafish Müller glia in contrast to the persistent reactive phase of Müller glia that does not result in cycle progression (Hoang et al., 2020). For the first time, we have defined different gene expression networks accompanying this transition in mature Müller glia of zebrafish larvae. Comparatively, we identified an intermediate expression module between quiescent and proliferative glia defined by activation of *notch3, fabp7a, id1* and *metrn*. Notch signalling is induced in the injured zebrafish retina (Wan et al., 2012), and influences Müller glia proliferation in the chick retina (Ghai et al., 2010), with its inhibition inducing Müller glia cell cycle entry (Conner et al., 2014; Campbell et al., 2021). ID signalling is attenuated by notch (Wang et al., 2009), and *id1* in Müller glia is associated with an inflammatory cell state (Todd et al., 2020). Also associated with these cells is *fabp7a*, with FABP-encoding genes selectively expressed in activated chick Müller glia (Hoang et al., 2020; Campbell et al., 2022). Finally, *metrn* is a glial differentiation gene that is associated with astrocyte gliosis (Nishino et al., 2004; Obayashi et al., 2009; Lee et al., 2015) but can also improve mammalian Müller glia cell cycle entry (Wang et al., 2012), as well as muscle regeneration (Lee et al., 2010;Baht et al., 2020). Therefore, despite genes in this transitioning module being implicated in mammalian gliosis, their upregulation in zebrafish Müller glia prior to cell cycle entry is likely important for proliferation.

These findings enhance our understanding of heterogeneous states of quiescent Müller glia and define the temporal sequence of gene expression upregulation and importantly downregulation to progress towards neurogenesis. Certain molecularly distinct states of Müller glia appear to influence whether a particular subset of Müller glia are primed for activation, with extrinsic cues building on from this state to induce Müller glia proliferation. Therefore, an understanding of the complex heterogeneity of resting-state glia will be critical when designing a blueprint towards improving mammalian Müller glia-driven, neural regeneration.

## 5 Conflict of Interest

The authors declare that the research was conducted in the absence of any commercial or financial relationships that could be construed as a potential conflict of interest.

## 6 Author Contributions

AK, SY, JH and PRJ conceived of and designed the study. AK and SY carried out most of the experiments and data analysis. AN performed pseudotime trajectory analysis. AK wrote the first draft and all authors revised the manuscript.

## 7 Funding

This study was supported by funding through a K.M. Brutton Bequest (University of Melbourne) and Research Grant Support Scheme (University of Melbourne). AK was supported by the Australian Government Research Training Program Scholarship.

## 8 Acknowledgments

We thank Dr. Ohshima and Prof. Wong for the plasmids they kindly provided. We acknowledge the staff of the Danio rerio University of Melbourne and Walter and Eliza Hall of Medical Institute fish facilities for animal husbandry and support. We acknowledge the support of the Biological Optical Microscopy Platform at the University of Melbourne. Manuscript editor Julian Heng (Remotely Consulting, Australia) provided professional English-language editing of this article.

---

**Supplementary Figure 1:***id1, fabp7a, notch3* and *her12* expression in Müller glia along pseudotime in integrated Lws2 and Sws2 photoreceptor ablation scRNAseq datasets.

**Supplementary Figure 2:** Expression of prominent markers upregulated before or during Müller glia proliferation.

**Supplementary Figure 3:** (A, C, E, G) Labelling for mCherry antibody reveals presence of photoreceptor debris (red) in Gfap-expressing Müller glia (grey) across injury models from 24-96 hours post injury (hpi). Scale bar = 50 μm. (B, D, F, H) Percentages of Müller glia containing mCherry photoreceptor debris. Cell counts represented as percentages of debris-containing Müller glia between activated and total glia cohorts following widespread Lws2 (I), reduced Lws2 (J) and Xops (K) ablation paradigms. * = p ≤ 0.05.

**Supplementary Figure 4:** (A) Enrichment term analysis of the Müller glia cluster expressing markers associated with stress response. (B) Heatmap visualizing the top 10 expressed genes of each quiescent Müller glia cluster (C1-C6).

**Supplementary Figure 5:** (A) RNAscope *in situ* hybridization of markers *fgf24* (A’), *efnb2a* (A”) and *rdh10a* (A”’) in the 12 month-old zebrafish retina.

**Supplementary Figure 6:** (A) Labelling of DAPI-positive nuclei (cyan), Gfap-expressing Müller glia and Bmpr1ba/b. Astrices indicate cell bodies of Müller glia in dorsal, central and ventral regions of the retina. Scale Bar = 10 μm. (B) Expression plots of the Bmpr1ba/b encoding genes.

**Supplementary Figure 7:** (A) Scatterplot indicating genes that are upregulated in resting Müller glia as a result of Lws2 ablation (photoreceptor ablation; pink) or reduced compared to Müller glia in uninjured (no ablation) retinas (cyan). Gene expression values are presented as log1p() expression values, averaged from all cells within the four quiescent Müller glia clusters highlighted in Figure 3. (B) Feature expression plots of genes *cxcl18b, cxcl14*, and *cd99*, in Müller glia from the uninjured (no ablation) and Lws2 ablation (photoreceptor ablation) datasets, split by condition.

## Notes

### Competing Interest Statement

The authors have declared no competing interest.

## References

Anderson, R.G.W., Kamen, B.A., Rothberg, K.G., and Lacey, S.W. (1992). Potocytosis: Sequestration and Transport of Small Molecules by Caveolae. Science 255, 410–411.

Ang, N.B., Saera-Vila, A., Walsh, C., Hitchcock, P.F., Kahana, A., Thummel, R., and Nagashima, M. (2020). Midkine-a functions as a universal regulator of proliferation during epimorphic regeneration in adult zebrafish. PLOS ONE 15, e0232308.

Atkinson-Leadbeater, K., Hehr, C.L., and Mcfarlane, S. (2014). Fgfr signaling is required as the early eye field forms to promote later patterning and morphogenesis of the eye. Developmental Dynamics 243, 663–675.

Baht, G.S., Bareja, A., Lee, D.E., Rao, R.R., Huang, R., Huebner, J.L., Bartlett, D.B., Hart, C.R., Gibson, J.R., Lanza, I.R., Kraus, V.B., Gregory, S.G., Spiegelman, B.M., and White, J.P. (2020). Meteorin-like facilitates skeletal muscle repair through a Stat3/IGF-1 mechanism. Nat Metab 2, 278–289.

Bailey, T.J., Fossum, S.L., Fimbel, S.M., Montgomery, J.E., and Hyde, D.R. (2010). The inhibitor of phagocytosis, O-phospho-L-serine, suppresses Muller glia proliferation and cone cell regeneration in the light-damaged zebrafish retina. Exp Eye Res 91, 601–612.

Baumgartner, W.A. (2000). Etiology, pathogenesis, and experimental treatment of retinitis pigmentosa. Medical Hypotheses 54, 814–824.

Bernardos, R.L., Barthel, L.K., Meyers, J.R., and Raymond, P.A. (2007). Late-Stage Neuronal Progenitors in the Retina Are Radial Müller Glia That Function as Retinal Stem Cells. The Journal of Neuroscience 27, 7028–7040.

Bernardos, R.L., and Raymond, P.A. (2006). GFAP transgenic zebrafish. Gene Expression Patterns 6, 1007–1013.

Blum, N., and Begemann, G. (2012). Retinoic acid signaling controls the formation, proliferation and survival of the blastema during adult zebrafish fin regeneration. Development 139, 107–116.

Bond, A.M., Bhalala, O.G., and Kessler, J.A. (2012). The dynamic role of bone morphogenetic proteins in neural stem cell fate and maturation. Dev Neurobiol 72, 1068–1084.

Bourne, R.R.A., Flaxman, S.R., Braithwaite, T., Cicinelli, M.V., Das, A., Jonas, J.B., Keeffe, J., Kempen, J.H., Leasher, J., Limburg, H., Naidoo, K., Pesudovs, K., Resnikoff, S., Silvester, A., Stevens, G.A., Tahhan, N., Wong, T.Y., Taylor, H.R., Bourne, R., Ackland, P., Arditi, A., Barkana, Y., Bozkurt, B., Braithwaite, T., Bron, A., Budenz, D., Cai, F., Casson, R., Chakravarthy, U., Choi, J., Cicinelli, M.V., Congdon, N., Dana, R., Dandona, R., Dandona, L., Das, A., Dekaris, I., Del Monte, M., Deva, J., Dreer, L., Ellwein, L., Frazier, M., Frick, K., Friedman, D., Furtado, J., Gao, H., Gazzard, G., George, R., Gichuhi, S., Gonzalez, V., Hammond, B., Hartnett, M.E., He, M., Hejtmancik, J., Hirai, F., Huang, J., Ingram, A., Javitt, J., Jonas, J., Joslin, C., Keeffe, J., Kempen, J., Khairallah, M., Khanna, R., Kim, J., Lambrou, G., Lansingh, V.C., Lanzetta, P., Leasher, J., Lim, J., Limburg, H., Mansouri, K., Mathew, A., Morse, A., Munoz, B., Musch, D., Naidoo, K., Nangia, V., Palaiou, M., Parodi, M.B., Pena, F.Y., Pesudovs, K., Peto, T., Quigley, H., Raju, M., Ramulu, P., Resnikoff, S., Robin, A., Rossetti, L., Saaddine, J., Sandar, M.Y.A., Serle, J., Shen, T., Shetty, R., Sieving, P., Silva, J.C., Silvester, A., Sitorus, R.S., Stambolian, D., Stevens, G., et al. (2017). Magnitude, temporal trends, and projections of the global prevalence of blindness and distance and near vision impairment: a systematic review and meta-analysis. The Lancet Global Health 5, e888–e897.

Boyd, P., and Hyde, D.R. (2022). Iron contributes to photoreceptor degeneration and Müller glia proliferation in the zebrafish light-treated retina. Experimental Eye Research 216, 108947.

Bradley, J. (2008). TNF-mediated inflammatory disease. The Journal of Pathology 214, 149–160.

Bringmann, A., Iandiev, I., Pannicke, T., Wurm, A., Hollborn, M., Wiedemann, P., Osborne, N.N., and Reichenbach, A. (2009a). Cellular signaling and factors involved in Müller cell gliosis: Neuroprotective and detrimental effects. Progress in Retinal and Eye Research 28, 423–451.

Bringmann, A., Pannicke, T., Biedermann, B., Francke, M., Iandiev, I., Grosche, J., Wiedemann, P., Albrecht, J., and Reichenbach, A. (2009b). Role of retinal glial cells in neurotransmitter uptake and metabolism. Neurochemistry International 54, 143–160.

Bringmann, A., Pannicke, T., Grosche, J., Francke, M., Wiedemann, P., Skatchkov, S.N., Osborne, N.N., and Reichenbach, A. (2006). Müller cells in the healthy and diseased retina. Progress in Retinal and Eye Research 25, 397–424.

Bringmann, A., and Wiedemann, P. (2012). Müller Glial Cells in Retinal Disease. Ophthalmologica 227, 1–19.

Campbell, L.J., Hobgood, J.S., Jia, M., Boyd, P., Hipp, R.I., and Hyde, D.R. (2021). Notch3 and DeltaB maintain Müller glia quiescence and act as negative regulators of regeneration in the light-damaged zebrafish retina. Glia 69, 546–566.

Campbell, W.A., Tangeman, A., El-Hodiri, H.M., Hawthorn, E.C., Hathoot, M., Blum, S., Hoang, T., Blackshaw, S., and Fischer, A.J. (2022). Fatty acid-binding proteins and fatty acid synthase influence glial reactivity and promote the formation of Müller glia-derived progenitor cells in the chick retina. Development 149.

Centanin, L., Ander, J.J., Hoeckendorf, B., Lust, K., Kellner, T., Kraemer, I., Urbany, C., Hasel, E., Harris, W.A., Simons, B.D., and Wittbrodt, J. (2014). Exclusive multipotency and preferential asymmetric divisions in post-embryonic neural stem cells of the fish retina. Development 141, 3472–3482.

Centanin, L., Hoeckendorf, B., and Wittbrodt, J. (2011). Fate restriction and multipotency in retinal stem cells. Cell Stem Cell 9, 553–562.

Christofori, G. (2003). Split personalities: the agonistic antagonist Sprouty. Nature Cell Biology 5, 377–379.

Close, J.L., Liu, J., Gumuscu, B., and Reh, T.A. (2006). Epidermal growth factor receptor expression regulates proliferation in the postnatal rat retina. Glia 54, 94–104.

Conedera, F.M., Pousa, A.M.Q., Mercader, N., Tschopp, M., and Enzmann, V. (2019). Retinal microglia signaling affects Müller cell behavior in the zebrafish following laser injury induction. Glia 67, 1150–1166.

Conedera, F.M., Pousa, A.M.Q., Mercader, N., Tschopp, M., and Enzmann, V. (2021). The TGFβ/Notch axis facilitates Müller cell-to-epithelial transition to ultimately form a chronic glial scar. Mol Neurodegener 16, 69.

Conner, C., Ackerman, K.M., Lahne, M., Hobgood, J.S., and Hyde, D.R. (2014). Repressing notch signaling and expressing TNFα are sufficient to mimic retinal regeneration by inducing Müller glial proliferation to generate committed progenitor cells. The Journal of neuroscience: the official journal of the Society for Neuroscience 34, 14403–14419.

Corradini, E., Babitt, J.L., and Lin, H.Y. (2009). The RGM/DRAGON family of BMP co-receptors. Cytokine & Growth Factor Reviews 20, 389–398.

Curado, S., Anderson, R.M., Jungblut, B., Mumm, J., Schroeter, E., and Stainier, D.Y.R. (2007). Conditional targeted cell ablation in zebrafish: A new tool for regeneration studies. Developmental Dynamics 236, 1025–1035.

Curado, S., Stainier, D.Y.R., and Anderson, R.M. (2008). Nitroreductase-mediated cell/tissue ablation in zebrafish: a spatially and temporally controlled ablation method with applications in developmental and regeneration studies. Nature protocols 3, 948–954.

D’orazi, F.D., Suzuki, S.C., Darling, N., Wong, R.O., and Yoshimatsu, T. (2020). Conditional and biased regeneration of cone photoreceptor types in the zebrafish retina. Journal of Comparative Neurology 528, 2816–2830.

Da Silva, S., and Cepko, C.L. (2017). Fgf8 Expression and Degradation of Retinoic Acid Are Required for Patterning a High-Acuity Area in the Retina. Dev Cell 42, 68–81.e66.

De La Rosa-Reyes, V., Duprey-Díaz, M.V., Blagburn, J.M., and Blanco, R.E. (2021). Retinoic acid treatment recruits macrophages and increases axonal regeneration after optic nerve injury in the frog Rana pipiens. PLoS One 16, e0255196.

Di Guglielmo, G.M., Le Roy, C., Goodfellow, A.F., and Wrana, J.L. (2003). Distinct endocytic pathways regulate TGF-β receptor signalling and turnover. Nature Cell Biology 5, 410–421.

Duprey-Díaz, M.V., Blagburn, J.M., and Blanco, R.E. (2016). Optic nerve injury upregulates retinoic acid signaling in the adult frog visual system. J Chem Neuroanat 77, 80–92.

Dyer, M.A., and Cepko, C.L. (2000). Control of Müller glial cell proliferation and activation following retinal injury. Nature Neuroscience 3, 873–880.

Elsaeidi, F., Macpherson, P., Mills, E.A., Jui, J., Flannery, J.G., and Goldman, D. (2018). Notch Suppression Collaborates with Ascl1 and Lin28 to Unleash a Regenerative Response in Fish Retina, But Not in Mice. The Journal of neuroscience: the official journal of the Society for Neuroscience 38, 2246–2261.

Fausett, B.V., and Goldman, D. (2006). A role for alpha1 tubulin-expressing Müller glia in regeneration of the injured zebrafish retina. The Journal of neuroscience: the official journal of the Society for Neuroscience 26, 6303–6313.

Fausett, B.V., Gumerson, J.D., and Goldman, D. (2008). The Proneural Basic Helix-Loop-Helix Gene <em>Ascl1a</em> Is Required for Retina Regeneration. The Journal of Neuroscience 28, 1109–1117.

Felfly, H., and Klein, O.D. (2013). Sprouty genes regulate proliferation and survival of human embryonic stem cells. Scientific Reports 3, 2277.

Ferrari, S., Di Iorio, E., Barbaro, V., Ponzin, D., Sorrentino, F.S., and Parmeggiani, F. (2011). Retinitis pigmentosa: genes and disease mechanisms. Current genomics 12, 238–249.

Fischer, A.J., Bosse, J.L., and El-Hodiri, H.M. (2013). The ciliary marginal zone (CMZ) in development and regeneration of the vertebrate eye. Experimental Eye Research 116, 199–204.

Fischer, A.J., Mcguire, C.R., Dierks, B.D., and Reh, T.A. (2002). Insulin and fibroblast growth factor 2 activate a neurogenic program in Müller glia of the chicken retina. J Neurosci 22, 9387–9398.

Fischer, A.J., Zelinka, C., Gallina, D., Scott, M.A., and Todd, L. (2014). Reactive microglia and macrophage facilitate the formation of Müller glia-derived retinal progenitors. Glia 62, 1608–1628.

Fleckenstein, M., Keenan, T.D.L., Guymer, R.H., Chakravarthy, U., Schmitz-Valckenberg, S., Klaver, C.C., Wong, W.T., and Chew, E.Y. (2021). Age-related macular degeneration. Nature Reviews Disease Primers 7, 31.

Furuya, T., Pan, Z., and Kashiwagi, K. (2012). Role of Retinal Glial Cell Glutamate Transporters in Retinal Ganglion Cell Survival Following Stimulation of NMDA Receptor. Current Eye Research 37, 170–178.

Gallina, D., Zelinka, C., and Fischer, A.J. (2014). Glucocorticoid receptors in the retina, Müller glia and the formation of Müller glia-derived progenitors. Development 141, 3340–3351.

Ghai, K., Zelinka, C., and Fischer, A.J. (2010). Notch signaling influences neuroprotective and proliferative properties of mature Müller glia. The Journal of neuroscience: the official journal of the Society for Neuroscience 30, 3101–3112.

Goldshmit, Y., Tang, J., Siegel, A.L., Nguyen, P.D., Kaslin, J., Currie, P.D., and Jusuf, P.R. (2018). Different Fgfs have distinct roles in regulating neurogenesis after spinal cord injury in zebrafish. Neural Dev 13, 24.

Gordois, A., Cutler, H., Pezzullo, L., Gordon, K., Cruess, A., Winyard, S., Hamilton, W., and Chua, K. (2012). An estimation of the worldwide economic and health burden of visual impairment. Global Public Health 7, 465–481.

Gorsuch, R.A., and Hyde, D.R. (2014). Regulation of Müller glial dependent neuronal regeneration in the damaged adult zebrafish retina. Experimental eye research 123, 131–140.

Hamel, C. (2006). Retinitis pigmentosa. Orphanet J Rare Dis 1, 40.

Hamon, A., García-García, D., Ail, D., Bitard, J., Chesneau, A., Dalkara, D., Locker, M., Roger, J.E., and Perron, M. (2019). Linking YAP to Müller Glia Quiescence Exit in the Degenerative Retina. Cell Reports 27, 1712–1725.e1716.

Harada, T., Harada, C., Kohsaka, S., Wada, E., Yoshida, K., Ohno, S., Mamada, H., Tanaka, K., Parada, L.F., and Wada, K. (2002). Microglia–Müller Glia Cell Interactions Control Neurotrophic Factor Production during Light-Induced Retinal Degeneration. The Journal of Neuroscience 22, 9228–9236.

Hartung, A., Bitton-Worms, K., Rechtman, M.M., Wenzel, V., Boergermann, J.H., Hassel, S., Henis, Y.I., and Knaus, P. (2006). Different routes of bone morphogenic protein (BMP) receptor endocytosis influence BMP signaling. Mol Cell Biol 26, 7791–7805.

Hauck, S.M., Kinkl, N., Deeg, C.A., Swiatek-De Lange, M., Schöffmann, S., and Ueffing, M. (2006). GDNF family ligands trigger indirect neuroprotective signaling in retinal glial cells. Molecular and cellular biology 26, 2746–2757.

Hayashi, T., Asano, Y., Shintani, Y., Aoyama, H., Kioka, H., Tsukamoto, O., Hikita, M., Shinzawa-Itoh, K., Takafuji, K., Higo, S., Kato, H., Yamazaki, S., Matsuoka, K., Nakano, A., Asanuma, H., Asakura, M., Minamino, T., Goto, Y., Ogura, T., Kitakaze, M., Komuro, I., Sakata, Y., Tsukihara, T., Yoshikawa, S., and Takashima, S. (2015). Higd1a is a positive regulator of cytochrome c oxidase. Proc Natl Acad Sci U S A 112, 1553–1558.

Heldin, C.-H., Landström, M., and Moustakas, A. (2009). Mechanism of TGF-β signaling to growth arrest, apoptosis, and epithelial–mesenchymal transition. Current Opinion in Cell Biology 21, 166–176.

Hoang, T., Wang, J., Boyd, P., Wang, F., Santiago, C., Jiang, L., Yoo, S., Lahne, M., Todd, L.J., Jia, M., Saez, C., Keuthan, C., Palazzo, I., Squires, N., Campbell, W.A., Rajaii, F., Parayil, T., Trinh, V., Kim, D.W., Wang, G., Campbell, L.J., Ash, J., Fischer, A.J., Hyde, D.R., Qian, J., and Blackshaw, S. (2020). Gene regulatory networks controlling vertebrate retinal regeneration. Science 370, eabb8598.

Hui, S.P., Sengupta, D., Lee, S.G.P., Sen, T., Kundu, S., Mathavan, S., and Ghosh, S. (2014). Genome Wide Expression Profiling during Spinal Cord Regeneration Identifies Comprehensive Cellular Responses in Zebrafish. PLOS ONE 9, e84212.

Iribarne, M., Hyde, D.R., and Masai, I. (2019). TNFα Induces Müller Glia to Transition From Non-proliferative Gliosis to a Regenerative Response in Mutant Zebrafish Presenting Chronic Photoreceptor Degeneration. Frontiers in Cell and Developmental Biology 7.

Itoh, S., and Ten Dijke, P. (2007). Negative regulation of TGF-β receptor/Smad signal transduction. Current Opinion in Cell Biology 19, 176–184.

Jorstad, N.L., Wilken, M.S., Grimes, W.N., Wohl, S.G., Vandenbosch, L.S., Yoshimatsu, T., Wong, R.O., Rieke, F., and Reh, T.A. (2017). Stimulation of functional neuronal regeneration from Müller glia in adult mice. Nature 548, 103–107.

Jorstad, N.L., Wilken, M.S., Todd, L., Finkbeiner, C., Nakamura, P., Radulovich, N., Hooper, M.J., Chitsazan, A., Wilkerson, B.A., Rieke, F., and Reh, T.A. (2020). STAT Signaling Modifies Ascl1 Chromatin Binding and Limits Neural Regeneration from Muller Glia in Adult Mouse Retina. Cell Reports 30, 2195–2208.e2195.

Kalloniatis, M., and Fletcher, E.L. (2004). Retinitis pigmentosa: understanding the clinical presentation, mechanisms and treatment options. Clinical and Experimental Optometry 87, 65–80.

Karl, M.O., Hayes, S., Nelson, B.R., Tan, K., Buckingham, B., and Reh, T.A. (2008). Stimulation of neural regeneration in the mouse retina. Proc Natl Acad Sci U S A 105, 19508–19513.

Kase, S., Yoshida, K., Harada, T., Harada, C., Namekata, K., Suzuki, Y., Ohgami, K., Shiratori, K., Nakayama, K.I., and Ohno, S. (2006). Phosphorylation of extracellular signal-regulated kinase and p27(KIP1) after retinal detachment. Graefe’s Archive for Clinical and Experimental Ophthalmology 244, 352–358.

Kent, M.R., Kara, N., and Patton, J.G. (2021). Inhibition of GABA(A)-ρ receptors induces retina regeneration in zebrafish. Neural Regen Res 16, 367–374.

Ko, C.-Y., Chang, W.-C., and Wang, J.-M. (2015). Biological roles of CCAAT/Enhancer-binding protein delta during inflammation. Journal of Biomedical Science 22, 6.

Kuracha, M.R., Burgess, D., Siefker, E., Cooper, J.T., Licht, J.D., Robinson, M.L., and Govindarajan, V. (2011). Spry1 and Spry2 Are Necessary for Lens Vesicle Separation and Corneal Differentiation. Investigative Ophthalmology & Visual Science 52, 6887–6897.

Lahne, M., Brecker, M., Jones, S.E., and Hyde, D.R. (2021). The Regenerating Adult Zebrafish Retina Recapitulates Developmental Fate Specification Programs. Frontiers in Cell and Developmental Biology 8.

Lamba, D., Karl, M., and Reh, T. (2008). Neural regeneration and cell replacement: a view from the eye. Cell stem cell 2, 538–549.

Le, Y.Z. (2017). VEGF production and signaling in Müller glia are critical to modulating vascular function and neuronal integrity in diabetic retinopathy and hypoxic retinal vascular diseases. Vision Res 139, 108–114.

Leach, L.L., Hanovice, N.J., George, S.M., Gabriel, A.E., and Gross, J.M. (2021). The immune response is a critical regulator of zebrafish retinal pigment epithelium regeneration. Proceedings of the National Academy of Sciences 118, e2017198118.

Lee, H.S., Han, J., Lee, S.H., Park, J.A., and Kim, K.W. (2010). Meteorin promotes the formation of GFAP-positive glia via activation of the Jak-STAT3 pathway. J Cell Sci 123, 1959–1968.

Lee, H.S., Lee, S.H., Cha, J.H., Seo, J.H., Ahn, B.J., and Kim, K.W. (2015). Meteorin is upregulated in reactive astrocytes and functions as a negative feedback effector in reactive gliosis. Mol Med Rep 12, 1817–1823.

Lee, K.J., Kim, Y., Kim, M.S., Ju, H.M., Choi, B., Lee, H., Jeoung, D., Moon, K.W., Kang, D., Choi, J., Yook, J.I., and Hahn, J.H. (2020a). CD99-PTPN12 Axis Suppresses Actin Cytoskeleton-Mediated Dimerization of Epidermal Growth Factor Receptor. Cancers (Basel) 12.

Lee, M.-S., Wan, J., and Goldman, D. (2020b). Tgfb3 collaborates with PP2A and notch signaling pathways to inhibit retina regeneration. eLife 9, e55137.

Lenkowski, J.R., Qin, Z., Sifuentes, C.J., Thummel, R., Soto, C.M., Moens, C.B., and Raymond, P.A. (2013). Retinal regeneration in adult zebrafish requires regulation of TGFβ signaling. Glia 61, 1687–1697.

Lenkowski, J.R., and Raymond, P.A. (2014). Müller glia: Stem cells for generation and regeneration of retinal neurons in teleost fish. Progress in retinal and eye research 40, 94–123.

Lew, D.S., Mcgrath, M.J., and Finnemann, S.C. (2022). Galectin-3 Promotes Müller Glia Clearance Phagocytosis via MERTK and Reduces Harmful Müller Glia Activation in Inherited and Induced Retinal Degeneration. Frontiers in Cellular Neuroscience 16.

Lopez-Ramirez, M.A., Calvo, C.F., Ristori, E., Thomas, J.L., and Nicoli, S. (2016). Isolation and Culture of Adult Zebrafish Brain-derived Neurospheres. J Vis Exp, 53617.

Lourenço, R., Brandão, A.S., Borbinha, J., Gorgulho, R., and Jacinto, A. (2021). Yap Regulates Müller Glia Reprogramming in Damaged Zebrafish Retinas. Front Cell Dev Biol 9, 667796.

Lu, J., Chatterjee, M., Schmid, H., Beck, S., and Gawaz, M. (2016). CXCL14 as an emerging immune and inflammatory modulator. Journal of Inflammation 13, 1.

Macdonald, R.B., Randlett, O., Oswald, J., Yoshimatsu, T., Franze, K., and Harris, W.A. (2015). Müller glia provide essential tensile strength to the developing retina. The Journal of cell biology 210, 1075–1083.

Masland, R.H. (2012). The neuronal organization of the retina. Neuron 76, 266–280.

Medrano, M.P., Bejarano, C.A., Battista, A.G., Venera, G.D., Bernabeu, R.O., and Faillace, M.P. (2017). Injury-induced purinergic signalling molecules upregulate pluripotency gene expression and mitotic activity of progenitor cells in the zebrafish retina. Purinergic signalling 13, 443–465.

Mitchell, D.M., Sun, C., Hunter, S.S., New, D.D., and Stenkamp, D.L. (2019). Regeneration associated transcriptional signature of retinal microglia and macrophages. Scientific Reports 9, 4768.

Mitra, S., Sharma, P., Kaur, S., Khursheed, M.A., Gupta, S., Ahuja, R., Kurup, A.J., Chaudhary, M., and Ramachandran, R. (2018). Histone Deacetylase-Mediated Müller Glia Reprogramming through Her4.1-Lin28a Axis Is Essential for Retina Regeneration in Zebrafish. iScience 7, 68–84.

Montgomery, J.E., Parsons, M.J., and Hyde, D.R. (2010). A novel model of retinal ablation demonstrates that the extent of rod cell death regulates the origin of the regenerated zebrafish rod photoreceptors. The Journal of comparative neurology 518, 800–814.

Nagashima, M., Barthel, L.K., and Raymond, P.A. (2013). A self-renewing division of zebrafish Müller glial cells generates neuronal progenitors that require N-cadherin to regenerate retinal neurons. Development (Cambridge, England) 140, 4510–4521.

Nelson, C.M., Ackerman, K.M., O’hayer, P., Bailey, T.J., Gorsuch, R.A., and Hyde, D.R. (2013). Tumor necrosis factor-alpha is produced by dying retinal neurons and is required for Muller glia proliferation during zebrafish retinal regeneration. The Journal of neuroscience: the official journal of the Society for Neuroscience 33, 6524–6539.

Newman, E.A., Frambach, D.A., and Odette, L.L. (1984). Control of extracellular potassium levels by retinal glial cell K+ siphoning. Science (New York, N.Y.) 225, 1174–1175.

Ng Chi Kei, J., Currie, P.D., and Jusuf, P.R. (2017). Fate bias during neural regeneration adjusts dynamically without recapitulating developmental fate progression. Neural Development 12, 12.

Ng, J., Currie, P.D., and Jusuf, P.R. (2014). “The Regenerative Potential of the Vertebrate Retina: Lessons from the Zebrafish,” in Regenerative Biology of the Eye, ed. A. Pébay. (New York, NY: Springer New York), 49–82.

Nishino, J., Yamashita, K., Hashiguchi, H., Fujii, H., Shimazaki, T., and Hamada, H. (2004). Meteorin: a secreted protein that regulates glial cell differentiation and promotes axonal extension. Embo j 23, 1998–2008.

Noel, N.C.L., Macdonald, I.M., and Allison, W.T. (2021). Zebrafish Models of Photoreceptor Dysfunction and Degeneration. Biomolecules 11.

Nomura-Komoike, K., Saitoh, F., and Fujieda, H. (2020). Phosphatidylserine recognition and Rac1 activation are required for Muller glia proliferation, gliosis and phagocytosis after retinal injury. Sci Rep 10, 1488.

Obayashi, S., Tabunoki, H., Kim, S.U., and Satoh, J. (2009). Gene expression profiling of human neural progenitor cells following the serum-induced astrocyte differentiation. Cell Mol Neurobiol 29, 423–438.

Pollak, J., Wilken, M.S., Ueki, Y., Cox, K.E., Sullivan, J.M., Taylor, R.J., Levine, E.M., and Reh, T.A. (2013). ASCL1 reprograms mouse Müller glia into neurogenic retinal progenitors. Development 140, 2619–2631.

Powell, C., Cornblath, E., Elsaeidi, F., Wan, J., and Goldman, D. (2016). Zebrafish Müller glia-derived progenitors are multipotent, exhibit proliferative biases and regenerate excess neurons. Scientific Reports 6, 24851.

Powell, C., Grant, A.R., Cornblath, E., and Goldman, D. (2013). Analysis of DNA methylation reveals a partial reprogramming of the Müller glia genome during retina regeneration. Proceedings of the National Academy of Sciences 110, 19814–19819.

Ramachandran, R., Fausett, B.V., and Goldman, D. (2010). Ascl1a regulates Müller glia dedifferentiation and retinal regeneration through a Lin-28-dependent, let-7 microRNA signalling pathway. Nature Cell Biology 12, 1101–1107.

Ramachandran, R., Zhao, X.-F., and Goldman, D. (2011). Ascl1a/Dkk/beta-catenin signaling pathway is necessary and glycogen synthase kinase-3beta inhibition is sufficient for zebrafish retina regeneration. Proceedings of the National Academy of Sciences of the United States of America 108, 15858–15863.

Ramachandran, R., Zhao, X.-F., and Goldman, D. (2012). Insm1a-mediated gene repression is essential for the formation and differentiation of Müller glia-derived progenitors in the injured retina. Nature cell biology 14, 1013–1023.

Rao, M.B., Didiano, D., and Patton, J.G. (2017). Neurotransmitter-Regulated Regeneration in the Zebrafish Retina. Stem cell reports 8, 831–842.

Reichenbach, A., and Bringmann, A. (2013). New functions of Müller cells. Glia 61, 651–678.

Reichenbach, A., and Bringmann, A. (2016). Purinergic signaling in retinal degeneration and regeneration. Neuropharmacology 104, 194–211.

Resta, V., Novelli, E., Vozzi, G., Scarpa, C., Caleo, M., Ahluwalia, A., Solini, A., Santini, E., Parisi, V., Di Virgilio, F., and Galli-Resta, L. (2007). Acute retinal ganglion cell injury caused by intraocular pressure spikes is mediated by endogenous extracellular ATP. European Journal of Neuroscience 25, 2741–2754.

Roesch, K., Stadler, M.B., and Cepko, C.L. (2012). Gene expression changes within Müller glial cells in retinitis pigmentosa. Mol Vis 18, 1197–1214.

Rueda, E.M., Hall, B.M., Hill, M.C., Swinton, P.G., Tong, X., Martin, J.F., and Poché, R.A. (2019). The Hippo Pathway Blocks Mammalian Retinal Müller Glial Cell Reprogramming. Cell reports 27, 1637–1649.e1636.

Sabin, K.Z., Jiang, P., Gearhart, M.D., Stewart, R., and Echeverri, K. (2019). AP-1(cFos/JunB)/miR-200a regulate the pro-regenerative glial cell response during axolotl spinal cord regeneration. Commun Biol 2, 91.

Sakami, S., Imanishi, Y., and Palczewski, K. (2019). Müller glia phagocytose dead photoreceptor cells in a mouse model of retinal degenerative disease. The FASEB Journal 33, 3680–3692.

Sakuta, H., Takahashi, H., Shintani, T., Etani, K., Aoshima, A., and Noda, M. (2006). Role of Bone Morphogenic Protein 2 in Retinal Patterning and Retinotectal Projection. The Journal of Neuroscience 26, 10868–10878.

Sarkar, A., Jayesh Sodha, S., Junnuthula, V., Kolimi, P., and Dyawanapelly, S. (2022). Novel and investigational therapies for wet and dry age-related macular degeneration. Drug Discovery Today 27, 2322–2332.

Schenkel, A.R., Mamdouh, Z., Chen, X., Liebman, R.M., and Muller, W.A. (2002). CD99 plays a major role in the migration of monocytes through endothelial junctions. Nature Immunology 3, 143–150.

Schreiber, M., Kolbus, A., Piu, F., Szabowski, A., Möhle-Steinlein, U., Tian, J., Karin, M., Angel, P., and Wagner, E.F. (1999). Control of cell cycle progression by c-Jun is p53 dependent. Genes Dev 13, 607–619.

Sen, J., Harpavat, S., Peters, M.A., and Cepko, C.L. (2005). Retinoic acid regulates the expression of dorsoventral topographic guidance molecules in the chick retina. Development 132, 5147–5159.

Sharma, P., Gupta, S., Chaudhary, M., Mitra, S., Chawla, B., Khursheed, M.A., Saran, N.K., and Ramachandran, R. (2020). Biphasic Role of Tgf-β Signaling during Müller Glia Reprogramming and Retinal Regeneration in Zebrafish. iScience 23, 100817–100817.

Shaulian, E., and Karin, M. (2001). AP-1 in cell proliferation and survival. Oncogene 20, 2390–2400.

Sherpa, T., Fimbel, S.M., Mallory, D.E., Maaswinkel, H., Spritzer, S.D., Sand, J.A., Li, L., Hyde, D.R., and Stenkamp, D.L. (2008). Ganglion cell regeneration following whole-retina destruction in zebrafish. Developmental neurobiology 68, 166–181.

Shimizu, Y., Ito, Y., Tanaka, H., and Ohshima, T. (2015). Radial glial cell-specific ablation in the adult Zebrafish brain. Genesis 53, 431–439.

Sommer, F., Torraca, V., and Meijer, A.H. (2020). Chemokine Receptors and Phagocyte Biology in Zebrafish. Frontiers in Immunology 11.

Spek, C.A., Aberson, H.L., Butler, J.M., De Vos, A.F., and Duitman, J. (2021). CEBPD Potentiates the Macrophage Inflammatory Response but CEBPD Knock-Out Macrophages Fail to Identify CEBPD-Dependent Pro-Inflammatory Transcriptional Programs. Cells 10.

Stevens, G.A., White, R.A., Flaxman, S.R., Price, H., Jonas, J.B., Keeffe, J., Leasher, J., Naidoo, K., Pesudovs, K., Resnikoff, S., Taylor, H., and Bourne, R.R.A. (2013). Global Prevalence of Vision Impairment and Blindness: Magnitude and Temporal Trends, 1990–2010. Ophthalmology 120, 2377–2384.

Tanaka, T. (2016). “Leukocyte Adhesion Molecules,” in Encyclopedia of Immunobiology, ed. M.J.H. Ratcliffe. (Oxford: Academic Press), 505–511.

Taniguchi, K., Ayada, T., Ichiyama, K., Kohno, R., Yonemitsu, Y., Minami, Y., Kikuchi, A., Maehara, Y., and Yoshimura, A. (2007). Sprouty2 and Sprouty4 are essential for embryonic morphogenesis and regulation of FGF signaling. Biochem Biophys Res Commun 352, 896–902.

Tappeiner, C., Maurer, E., Sallin, P., Bise, T., Enzmann, V., and Tschopp, M. (2016). Inhibition of the TGFβ Pathway Enhances Retinal Regeneration in Adult Zebrafish. PLoS One 11, e0167073.

Thomas, J.L., Morgan, G.W., Dolinski, K.M., and Thummel, R. (2018). Characterization of the pleiotropic roles of Sonic Hedgehog during retinal regeneration in adult zebrafish. Experimental eye research 166, 106–115.

Thomas, J.L., Ranski, A.H., Morgan, G.W., and Thummel, R. (2016). Reactive gliosis in the adult zebrafish retina. Experimental Eye Research 143, 98–109.

Todd, L., Finkbeiner, C., Wong, C.K., Hooper, M.J., and Reh, T.A. (2020). Microglia Suppress Ascl1-Induced Retinal Regeneration in Mice. Cell Reports 33, 108507.

Todd, L., Hooper, M.J., Haugan, A.K., Finkbeiner, C., Jorstad, N., Radulovich, N., Wong, C.K., Donaldson, P.C., Jenkins, W., Chen, Q., Rieke, F., and Reh, T.A. (2021). Efficient stimulation of retinal regeneration from Müller glia in adult mice using combinations of proneural bHLH transcription factors. Cell Rep 37, 109857.

Todd, L., Squires, N., Suarez, L., and Fischer, A.J. (2016). Jak/Stat signaling regulates the proliferation and neurogenic potential of Müller glia-derived progenitor cells in the avian retina. Scientific Reports 6, 35703.

Todd, L., Suarez, L., Quinn, C., and Fischer, A.J. (2018). Retinoic Acid-Signaling Regulates the Proliferative and Neurogenic Capacity of Müller Glia-Derived Progenitor Cells in the Avian Retina. Stem Cells 36, 392–405.

Torraca, V., Otto, N.A., Tavakoli-Tameh, A., and Meijer, A.H. (2017). The inflammatory chemokine Cxcl18b exerts neutrophil-specific chemotaxis via the promiscuous chemokine receptor Cxcr2 in zebrafish. Dev Comp Immunol 67, 57–65.

Tsata, V., Kroehne, V., Wehner, D., Rost, F., Lange, C., Hoppe, C., Kurth, T., Reinhardt, S., Petzold, A., Dahl, A., Loeffler, M., Reimer, M.M., and Brand, M. (2020). Reactive oligodendrocyte progenitor cells (re-)myelinate the regenerating zebrafish spinal cord. Development 147.

Tsujimura, T., Hosoya, T., and Kawamura, S. (2010). A single enhancer regulating the differential expression of duplicated red-sensitive opsin genes in zebrafish. PLoS Genet 6, e1001245.

Ueki, Y., Wilken, M.S., Cox, K.E., Chipman, L., Jorstad, N., Sternhagen, K., Simic, M., Ullom, K., Nakafuku, M., and Reh, T.A. (2015). Transgenic expression of the proneural transcription factor Ascl1 in Müller glia stimulates retinal regeneration in young mice. Proc Natl Acad Sci U S A 112, 13717–13722.

Vihtelic, T.S., and Hyde, D.R. (2000). Light-induced rod and cone cell death and regeneration in the adult albino zebrafish (Danio rerio) retina. J Neurobiol 44, 289–307.

Wan, J., and Goldman, D. (2017). Opposing Actions of Fgf8a on Notch Signaling Distinguish Two Muller Glial Cell Populations that Contribute to Retina Growth and Regeneration. Cell reports 19, 849–862.

Wan, J., Ramachandran, R., and Goldman, D. (2012). HB-EGF is necessary and sufficient for Müller glia dedifferentiation and retina regeneration. Developmental cell 22, 334–347.

Wan, J., Zhao, X.-F., Vojtek, A., and Goldman, D. (2014). Retinal injury, growth factors, and cytokines converge on β-catenin and pStat3 signaling to stimulate retina regeneration. Cell reports 9, 285–297.

Wang, H.C., Perry, S.S., and Sun, X.H. (2009). Id1 attenuates Notch signaling and impairs T-cell commitment by elevating Deltex1 expression. Mol Cell Biol 29, 4640–4652.

Wang, R.N., Green, J., Wang, Z., Deng, Y., Qiao, M., Peabody, M., Zhang, Q., Ye, J., Yan, Z., Denduluri, S., Idowu, O., Li, M., Shen, C., Hu, A., Haydon, R.C., Kang, R., Mok, J., Lee, M.J., Luu, H.L., and Shi, L.L. (2014). Bone Morphogenetic Protein (BMP) signaling in development and human diseases. Genes Dis 1, 87–105.

Wang, S.M., Hsu, J.C., Ko, C.Y., Chiu, N.E., Kan, W.M., Lai, M.D., and Wang, J.M. (2016). Astrocytic CCAAT/Enhancer-Binding Protein Delta Contributes to Glial Scar Formation and Impairs Functional Recovery After Spinal Cord Injury. Mol Neurobiol 53, 5912–5927.

Wang, Z., Andrade, N., Torp, M., Wattananit, S., Arvidsson, A., Kokaia, Z., Jørgensen, J.R., and Lindvall, O. (2012). Meteorin is a chemokinetic factor in neuroblast migration and promotes stroke-induced striatal neurogenesis. J Cereb Blood Flow Metab 32, 387–398.

White, D.T., Sengupta, S., Saxena, M.T., Xu, Q., Hanes, J., Ding, D., Ji, H., and Mumm, J.S. (2017). Immunomodulation-accelerated neuronal regeneration following selective rod photoreceptor cell ablation in the zebrafish retina. Proc Natl Acad Sci U S A 114, E3719–E3728.

Wilhelmsson, U., Li, L., Pekna, M., Berthold, C.-H., Blom, S., Eliasson, C., Renner, O., Bushong, E., Ellisman, M., Morgan, T.E., and Pekny, M. (2004). Absence of glial fibrillary acidic protein and vimentin prevents hypertrophy of astrocytic processes and improves post-traumatic regeneration. The Journal of neuroscience: the official journal of the Society for Neuroscience 24, 5016–5021.

Wu, Q., Sun, C.C., Lin, H.Y., and Babitt, J.L. (2012). Repulsive Guidance Molecule (RGM) Family Proteins Exhibit Differential Binding Kinetics for Bone Morphogenetic Proteins (BMPs). PLOS ONE 7, e46307.

Xiong, B., Rui, Y., Zhang, M., Shi, K., Jia, S., Tian, T., Yin, K., Huang, H., Lin, S., Zhao, X., Chen, Y., Chen, Y.-G., Lin, S.-C., and Meng, A. (2006). Tob1 Controls Dorsal Development of Zebrafish Embryos by Antagonizing Maternal &#x3b2;-Catenin Transcriptional Activity. Developmental Cell 11, 225–238.

Xu, B., Tang, X., Jin, M., Zhang, H., Du, L., Yu, S., and He, J. (2020). Unifying developmental programs for embryonic and postembryonic neurogenesis in the zebrafish retina. Development 147, dev185660.

Xu, M., Zhai, Y., and Macdonald, I.M. (2020). Visual Field Progression in Retinitis Pigmentosa. Investigative Ophthalmology & Visual Science 61, 56–56.

Yamada, H., Yamada, E., Hackett, S.F., Ozaki, H., Okamoto, N., and Campochiaro, P.A. (1999). Hyperoxia causes decreased expression of vascular endothelial growth factor and endothelial cell apoptosis in adult retina. Journal of Cellular Physiology 179, 149–156.

Yao, K., Qiu, S., Tian, L., Snider, W.D., Flannery, J.G., Schaffer, D.V., and Chen, B. (2016). Wnt Regulates Proliferation and Neurogenic Potential of Müller Glial Cells via a Lin28/let-7 miRNA-Dependent Pathway in Adult Mammalian Retinas. Cell reports 17, 165–178.

Yao, K., Qiu, S., Wang, Y.V., Park, S.J.H., Mohns, E.J., Mehta, B., Liu, X., Chang, B., Zenisek, D., Crair, M.C., Demb, J.B., and Chen, B. (2018). Restoration of vision after de novo genesis of rod photoreceptors in mammalian retinas. Nature 560, 484–488.

Yoshida, K., Kase, S., Nakayama, K., Nagahama, H., Harada, T., Ikeda, H., Harada, C., Imaki, J., Ohgami, K., Shiratori, K., Ohno, S., and Nakayama, K.I. (2004). Distribution of p27(KIP1), cyclin D1, and proliferating cell nuclear antigen after retinal detachment. Graefe’s Archive for Clinical and Experimental Ophthalmology 242, 437–441.

Yoshimatsu, T., D’orazi, F.D., Gamlin, C.R., Suzuki, S.C., Suli, A., Kimelman, D., Raible, D.W., and Wong, R.O. (2016). Presynaptic partner selection during retinal circuit reassembly varies with timing of neuronal regeneration in vivo. Nat Commun 7, 10590.

Zenz, R., Eferl, R., Scheinecker, C., Redlich, K., Smolen, J., Schonthaler, H.B., Kenner, L., Tschachler, E., and Wagner, E.F. (2008). Activator protein 1 (Fos/Jun) functions in inflammatory bone and skin disease. Arthritis Research & Therapy 10, 201.

Zhang, C., Chaturvedi, D., Jaggar, L., Magnuson, D., Lee, J.M., and Patel, T.B. (2005). Regulation of Vascular Smooth Muscle Cell Proliferation and Migration by Human Sprouty 2. Arteriosclerosis, Thrombosis, and Vascular Biology 25, 533–538.

Zhang, J., Jiang, Z., Liu, X., and Meng, A. (2016). Eph/ephrin signaling maintains the boundary of dorsal forerunner cell cluster during morphogenesis of the zebrafish embryonic left-right organizer. Development 143, 2603–2615.

